# Surface-immobilized fibronectin conformation influences synovial fluid adsorption and film formation

**DOI:** 10.1101/2025.04.27.650860

**Authors:** Syeda Tajin Ahmed, Diego R. Jaramillo Pinto, Lenka Vitkova, Ummay Honey, Warren Flores, Katelyn L. Lunny, Kaleb A. Cutter, Yidan Wen, Kevin De France, Roberto C. Andresen Eguiluz

## Abstract

The articular cartilage extracellular matrix (ECM) is a complex network of biomolecules that includes fibronectin (FN). FN acts as an extracellular glue, controlling the assembly of other macromolecular constituents to the ECM. However, how FN participates in the binding and retention of synovial fluid components, the natural lubricant of articulated joints, to form a wear-protecting and lubricating film has not been established. This study reports on the role of FN and its molecular conformation in mediating macromolecular assembly of synovial fluid ad-layers. FN films as precursor films on functionalized surfaces, a model of FN’s articular cartilage surface, adsorbed and retained different amounts of synovial fluid (SF) depending on FN conformation. FN conformational changes were induced by depositing FN from bulk solution at pH 7 (extended state) or at pH 4 (unfolded state) on self-assembled monolayers on gold-coated quartz crystals, followed by adsorption of diluted SF (25%) onto FN precursor films. Mass density, thin film compliance, surface morphologies, and the secondary and tertiary structures of adsorbed FN films reveal pH-induced differences. FN films deposited at pH 4 were thicker, more rigid, showed a more homogeneous morphology, and had altered *α*-helix and *β*-sheet content, compared to FN films deposited at pH 7. FN precursor films deposited at pH 7 adsorbed and retained more synovial fluid than those at pH 4, revealing the importance of FN conformation at the articular cartilage surface to bind and maintain a thin layer of synovial fluid constituents. This knowledge will enable a better understanding of the molecular interactions and synergies between the articular cartilage ECM components and SF.

**GRAPHICAL ABSTRACT:** 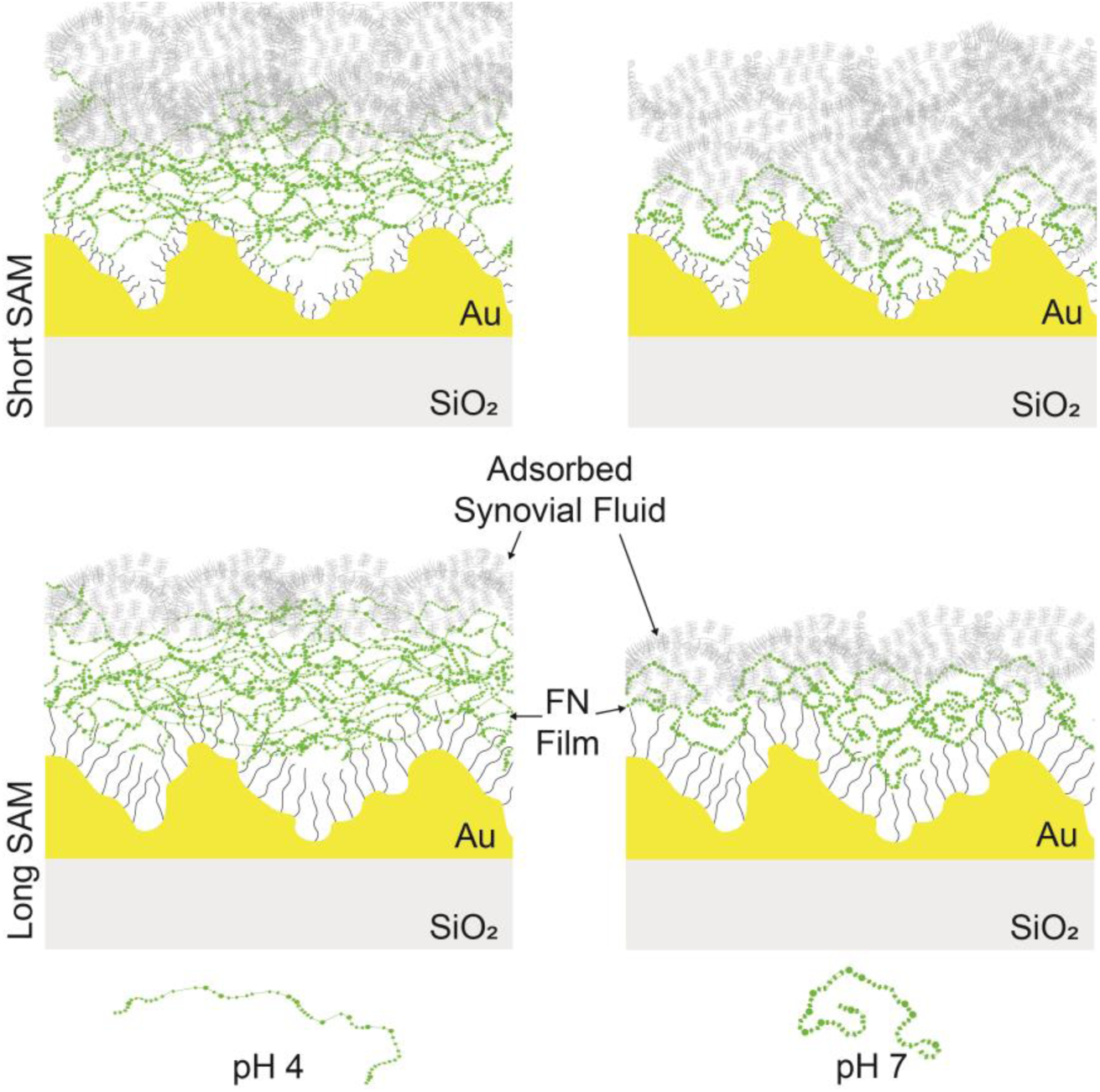

## INTRODUCTION

Fibronectin (FN) is a ⁓500 kDa dimeric glycoprotein, with each monomer consisting of 12 type I repeats (FN_I_), two type II repeats (FN_II_), and 15-17 type III (FN_III_) repeats.[1] FN_III_ repeats form *β*-barrels, and are more susceptible to conformational changes due to the lack of disulfide bonds than FN_I_ or FN_II_ repeats.[2] Its conformational flexibility makes it an essential component of the extracellular matrix (ECM) of tissues,[3,4] having a plethora of functions, including controlling cell adhesion and migration, being involved in the clotting cascade, and acting as a growth factor reservoir.[2,5,6] Its biological activity is dictated by its conformation, which can range from inactive in globular form (*e.g.*, soluble in blood plasma) to controlling binding partners or switching cellular behavior when insoluble, in extended, partially unfolded, or unfolded states (*e.g.*, when assembled into the ECM).[5,7] Furthermore, FN displays several surface-exposed molecular recognition sites for collagen, heparin, fibrin, among others.[1,2,8,9] Together, these binding sites provide FN with various biomolecular scaffolding functions. Additionally, several cryptic binding sites, sequences buried in the compact form of FN, become activated or deactivated due to conformational changes.[7,10,11]

The articular cartilage ECM is a complex network of biomolecules consisting mostly of heterotypic collagens, glycosaminoglycans, among others, including fibronectin [12–17]. Together, they contribute to articular joints’ biochemical and biomechanical homeostasis and proper function. Soluble FN in synovial fluid (SF), the natural lubricant of articulated joints, has been extensively studied: it has been identified as a marker for osteoarthritis [17–19], and FN fragments have been found to induce catabolic responses.[18–20] On the other hand, insoluble FN in the articular cartilage ECM, identified to increase in cartilage repair,[17] has been overlooked. FN in the articular cartilage ECM is found in a range of isoforms, with high levels of a variant containing the ED-B extra domain between FN_III-7_ and FN_III-8_ and a variant that lacks a region known as the V region, close to the C-terminal.[1,21] At the surface of articular cartilage ECM, FN is present and has been identified as a binding partner for SF components, including wear-protective and boundary-lubricating glycoproteins, such as lubricin.[22–24]

While the interactions between other interfacial ECM components, such as collagen II or hyaluronan, and SF molecules have been extensively investigated at the molecular scale in the context of synovial joint tribology,[25–31] few studies have addressed FN’s role in mediating SF component adsorption and retention, leading to the formation of a wear-protecting and lubricating film mediating cartilage tribology.[24,32] Building on previous studies and established protocols for the tunability of FN’s molecular conformation,[2,33,34] this study investigated molecular FN films formed from FN aqueous solutions at either a pH of 7.4 (extended FN molecular conformation) or a pH of 4 (partially unfolded/unfolded FN molecular conformation) on self-assembled monolayers (SAMs) and FN’s molecular film efficacy to adsorb and retain SF. Molecular FN film thickness, shear-dependent elastic compliance, film morphology, and molecular conformation were quantified using a combination of quartz crystal microbalance with dissipation (QCM-D), atomic force microscopy (AFM), and diffuse reflectance circular dichroism (DRCD).Their SF adsorption and retention capabilities were compared via QCM-D. Results indicate partially unfolded/unfolded FN (pH 4) molecules formed thicker films with altered *α*-helix content, less *β*-sheet content, and smoother morphologies than films formed from extended FN (pH 7) molecules. However, FN films formed from extended FN retained more SF than films formed from unfolded FN molecules. These molecular conformational changes lead to altered dSF film add layers.

Overall, this study presents a model to investigate the interaction between FN and SF, assisting in elucidating FN’s role in scaffolding a lubricating and wear-protecting film. The results demonstrate that FN conformation is involved in mediating the chemo-mechanical properties of the articular cartilage ECM and SF interface. This knowledge will enable a better understanding of the molecular regulation of articular cartilage-SF interface homeostasis and joint pathophysiology.

## EXPERIMENTAL SECTION

### Materials

Phosphate buffer saline (PBS) (Gibco, Cat. # 10-010-031), ethanol (ACROS, absolute, 200 Proof, ≥99.5%, Cat. # 61509-5000), sodium dodecyl sulfate (SDS) (MP Biomedicals, ultrapure, ≥99%, Cat. # 811032) were purchased and used as received unless otherwise indicated. Base-piranha solution was prepared with ammonium hydroxide (Chemsavers, Cat. # AMHE500ML, 28-30% pure), hydrogen peroxide (PERDROGEN^TM^ by Honeywell, MDL # MFCD00011333, ≥30% (w/w) stabilized). Gold-coated silica quartz crystals were purchased from Quartz PRO (Cat. # QCM5140CrAu120-050-Q, resonance frequency of 5 MHz). For gold surface functionalization, cysteamine hydrochloride (referred to from now on as short SAM) (Sigma, Cat. # M6500, ≥98% titration), 11-amino-1-undecanethiol hydrochloride (Dojindo, Cat. # 143339-58-6, ≥90% purity) (referred to from now on as long SAM), and glutaraldehyde (Sigma, 25% aqueous solution, Grade I, Cat. # G6257) were used. Ultrapure water was collected from a Thermo-Fisher Millipore UV water purification system. Bovine plasma fibronectin (FN) (Invitrogen, Cat. # 33010-018, 1 mg), and bovine synovial fluid (SF) (Lampire Biological Laboratories, Product # 8620853) were used for protein adsorption studies.

All experiments performed with FN and SF followed the procedures and guidelines provided by the University of California, Merced biosafety committee.

### Synovial fluid

Pooled bovine SF was used to prepare working aliquots, which were diluted to 25%v/v with PBS, kept frozen (*T* = −80 °C), and thawed on the day of the experiment. SF was first centrifuged for 5 min at 6000 rpm for all experiments to remove cell debris and large tissue aggregates. Next, the suspension was transferred to a new tube, where the fluid was diluted to 25% v/v to thin it for QCM-D measurements and prevent tubing clogging. This is referred to as dilute SF, or dSF in the text.

### Fibronectin (FN)

FN was diluted to a stock concentration of 1 mg/mL, kept frozen (*T* = −20 °C), and thawed on the day of the experiment. The working concentration of 50 µg/mL was prepared with PBS with a pH of 7.4, resulting in extended FN conformation in bulk solutions. PBS with a pH of 4 was prepared by adding tiny acetic acid droplets and was used within two weeks to induce FN conformational changes and unfold FN. [35]

### Functionalization of substrates with self-assembled monolayers for QCM-D

Gold-coated crystals of 5 MHz fundamental resonance frequency were rinsed copiously with ultrapure water (18.2 MΩ.cm), 2 wt% SDS, and ethanol, respectively, and repeated three times. After rinsing, surfaces were dried with a gentle stream of N_2_, followed by oxygen plasma (Harrick Plasma, PDC-32G-115V) for 2 minutes and stored in Petri dishes before use.

The clean crystals were incubated in 0.088M ethanolic solution of cysteamine hydrochloride for short SAMs or 0.042 M of 11-amino-1-undecanethiol hydrochloride for long SAMs at 4^°^C overnight. This process forms a self-assembled monolayer of NH_2_-terminated thiol on the gold surface. The next day, the surfaces were rinsed gently in ethanol to remove excess unbound NH_2_-terminated thiols and blown-dried with N_2_ gas. Then, the surfaces were incubated in 25% aqueous glutaraldehyde solution at room temperature for 30 minutes. This allows imine formation between the aldehyde group of glutaraldehyde and the amine group in the cysteamine chloride or 11-amino-1-undecanethiol hydrochloride, Figure S1. After incubation, the substrates were rinsed with water and then blown dry with a N_2_ stream. The functionalized surface was immediately loaded into the QCM-D fluidic cell.

After being used for protein adsorption experiments in the QCM-D, crystals were reused after cleaning in base piranha (6:1:1 by volume of water: ammonium hydroxide: hydrogen peroxide) by submerging the surfaces for ∼30 seconds at 60 °C. Then, the crystals were further rinsed with water and ethanol several times and dried with a gentle stream of dry N_2_, finishing with a short plasma cleaning step. Each crystal was reused on average 5 times.

### X-Ray Photoelectron Spectroscopy

A Thermoscientific Nexsa G2 was used to confirm the gold functionalization with amine-terminated SAMs (long and short) and estimate their relative densities. The same SAM functionalization procedure used to functionalize the QCM-D crystals was followed, without the glutaraldehyde step. Data was analyzed using the software KherveFitting v1.545. The binding energy was corrected by recording a C1s C-C peak for each sample and centering it at 284.6 eV. For the control bare Au crystal, Figure S2, the correction was done by adjusting the Au 4f_7/2_ peak to 84.0 eV due to the lack of C peaks. A total of 25 scans in a circular area with a diameter of 50 µm were performed on a point in the center of each sample, recording an overview spectrum (1 eV step size, 10 ms dwell time, 200 eV pass energy), and high-resolution spectra (0.1 eV step size, 50 ms dwell time, 50 eV pass energy) for C, S, N, Cl, O, and Au. The N 1s high-resolution spectra were used to estimate the difference in amine density between short and long SAMs, applying a Gaussian-Lorentzian fit to their counts per second (CPS) versus binding energy plots, calculating the area under the curves, and normalizing it by the area of the scanned region.

### Quartz Crystal Microbalance with Dissipation (QCM-D)

QCM-D measurements were performed using a QCM-I mini (Micro Vacuum, Hungary) and a Q-Sense Explorer microbalance (Biolin, Sweden), both with similar performance, using AT-cut quartz crystals with 5 MHz fundamental frequency coated with 50 nm of gold. All measurements were performed at 25 ^°^C, controlled by an incorporated Peltier unit. The outlet channel of the QCM-I mini and Q-Sense were connected to a peristaltic pump (BQ80S Microflow Variable-Speed Peristaltic Pump and ISMATEC, respectively), used to maintain a constant flow rate.

The QCM-D experimental sequence, shown in Figure 1, began with a PBS baseline at a pH of 7.4 (PBS 7) or a PBS solution with an adjusted pH of 4 (PBS 4), until a dispersion of 2 Hz/hr or less was achieved. Next, FN solutions of concentrations ranging from 10 to 100 µg/ml were forced to flow by applying suction using a peristaltic pump. After saturating the fluidic chamber, the peristaltic pump was stopped, and data were collected until a steady state was reached. This step is denoted as ‘FN pre-wash’ for the FN precursor film. PBS 7 or PBS 4 was used for rinsing after the deposition of FN in PBS 7 or PBS 4, respectively. This step is denoted as ‘FN post wash’ for the precursor film. For the experiments where precursor films were deposited at pH 4, a subsequent wash step was completed with PBS 7. Then, a blocking step with 10 mg/ml of bovine serum albumin (BSA) in PBS 7 was performed, followed by a wash with PBS 7. Finally, dSF was flowed and allowed to reach a steady state. This step is denoted as ‘dSF pre-wash’, finishing with a PBS 7 rinse, step denoted as ‘dSF post wash’.

**Figure 1.**
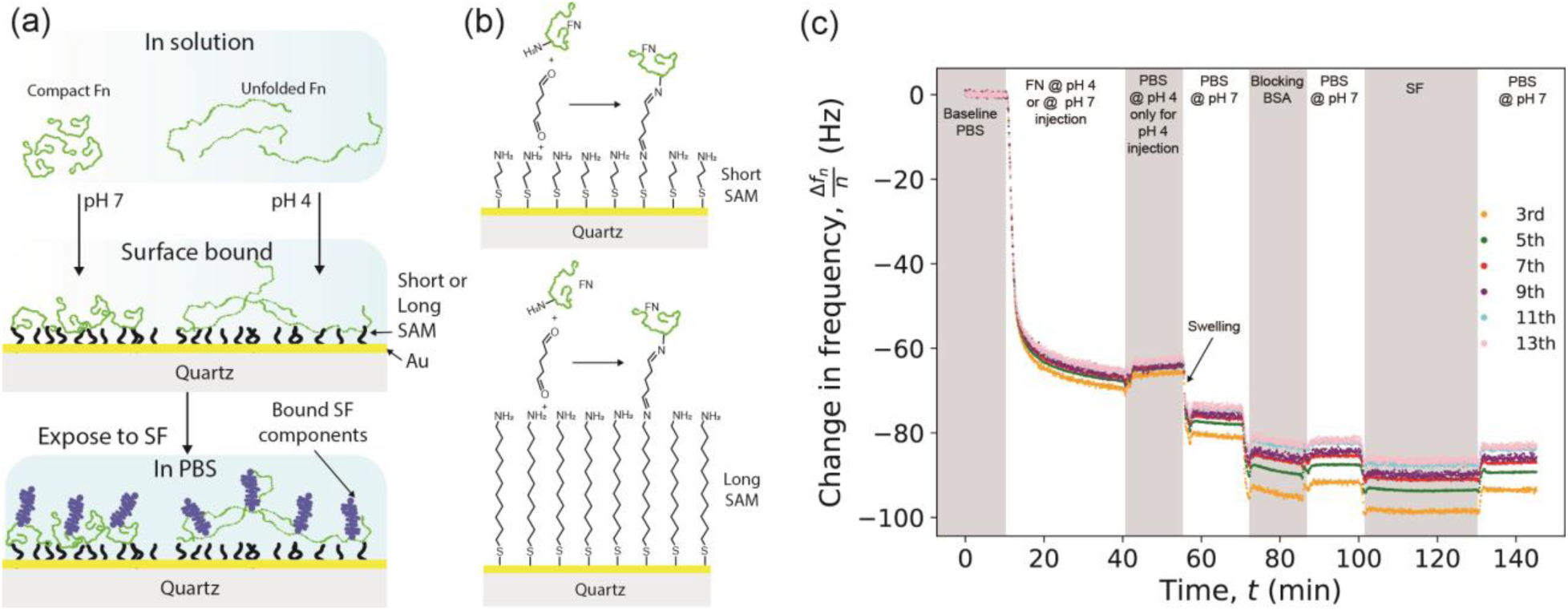
Schematics summarizing the experimental approach. (a) FN conformation was controlled in bulk solution by changing the pH. (b) Chemical immobilization of FN to short or long amine-terminated SAM surfaces with glutaraldehyde. (c) Change in frequency on a QCM-D sensor during a typical experiment.

The last 2-3 mins of the steady state values (Δ*f* and Δ*D*) of FN pre- and post-wash, as well as dSF pre- and post-wash, were used to calculate the average Sauerbrey mass, *m*_Sauerbrey_, the average Sauerbrey thickness, *d*_Sauerbrey_, and thin film elastic compliances, *J*^′^_f_, using the methods described below. Each reported condition consists of at least three independent measurements, reported as the average ± standard error of the mean, unless otherwise stated.

### Modeling of QCM-D data

Changes in frequency from the 3^rd^ to the 13^th^ overtones were used to compute the Sauerbrey mass using pyQCM-BRATADIO.[36] Briefly, the method uses linear regression to calculate the slope of the change in frequency as a function of the overtones, as illustrated in Figure S3a:

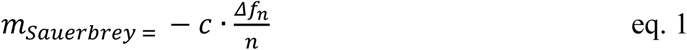

where *m*_Sauerbrey_ is the Sauerbrey mass in ng/cm^2^, *c* the crystal mass sensitivity, typically −17.7 ng·cm^-2^·Hz^-2^ for a 5 MHz crystal, and Δ*f*_n_ is the change in frequency for the overtone, *n*. The Sauerbrey thickness, *d*_Sauerbrey_, was estimated by assuming that the film density is similar to that of water, 1 g/cm³, and thus dividing *m*_Sauerbrey_ by a factor of 100.

The shear-dependent elastic compliance, *J*^′^_f_ (referred to from now on as elastic compliance) was computed using a thin film in liquids model, derived by Du and Johannsmann, as shown in Figure S3b:[37]

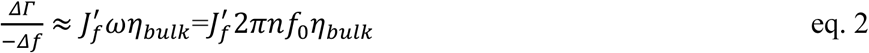

where *J*^′^_f_ is the elastic component of the shear elastic compliance of the thin film, and ΔΓ is the change in half-band half-width or the imaginary part of the resonant frequency. It is related to changes in dissipation *D* as follows:

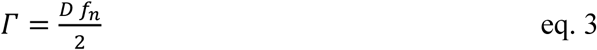

where *f*_n_ =*n*·*f*_o_ is the resonance frequency of the *n*^th^ overtone. Equation 2 can be applied to thin viscoelastic films deposited on a QCM-D crystal in liquid environments. It is assumed that the density of the bulk fluid and the adsorbed film are similar, *ρ*_bulk_ ≈ *ρ*_f_, where *ρ*_bulk_ and *ρ*_f_ are the densities of the bulk fluid and the adsorbed film, respectively.

QCM-D results are discussed around the Sauerbrey mass, *m*_Sauerbrey_, and elastic compliance, *J*^′^_f_, as their values are calculated with contributions from the frequency changes and dissipation changes of all available overtones. The Sauerbrey mass, *m*_Sauerbrey_, and thin film elastic compliance, *J*^′^_f_, of FN films formed on short and long SAM-functionalized Au surfaces were determined to be independent of FN bulk concentration tested in this study, between 10-100 µg/ml, Figure S4. As a result, the bulk concentration of FN at 50 µg/mL was chosen for the subsequent QCM-D, AFM, and DRCD analysis.

### Immobilization of FN for DRCD and AFM

Immobilization of FN for AFM and DRCD analysis was performed outside the QCM-D fluidic chamber as follows: a clean QCM-D crystal was functionalized with self-assembled monolayers (SAMs) and activated with glutaraldehyde, identical to the procedure used for QCM-D measurements. After the glutaraldehyde activation step, the crystal was rinsed with Milli-Q water, followed by complete immersion in a clean glass jar containing the 50 µg/mL FN solution at the corresponding pH for 2 hrs at 4 °C, ending with a PBS rinse.

### Diffuse Reflectance Circular Dichroism Spectroscopy (DRCD)

CD spectra were collected using a circular dichroism spectrometer (JASCO J-1500, Tokyo, Japan) equipped with an integrating sphere, operating in diffuse reflectance mode under a nitrogen atmosphere. Samples were placed into the sample holders and immersed in PBS at pH 7, as shown schematically in Figure 2. The sample holder was placed in the reflection measurement position. Spectra were collected as the average of 10 accumulations at a wavelength range of 340 nm to 190 nm, using a digital integration time of 4 seconds, bandwidth of 4 nm, and scanning speed of 100 nm/min. QCM sensors with short or long SAMs without any protein adsorbed were used as the baseline. The data was baseline corrected, zero-endpoint corrected, and smoothed using a Savitzky-Golay filter [38] with polynomial order 2 and frame length 9.

**Figure 2.**
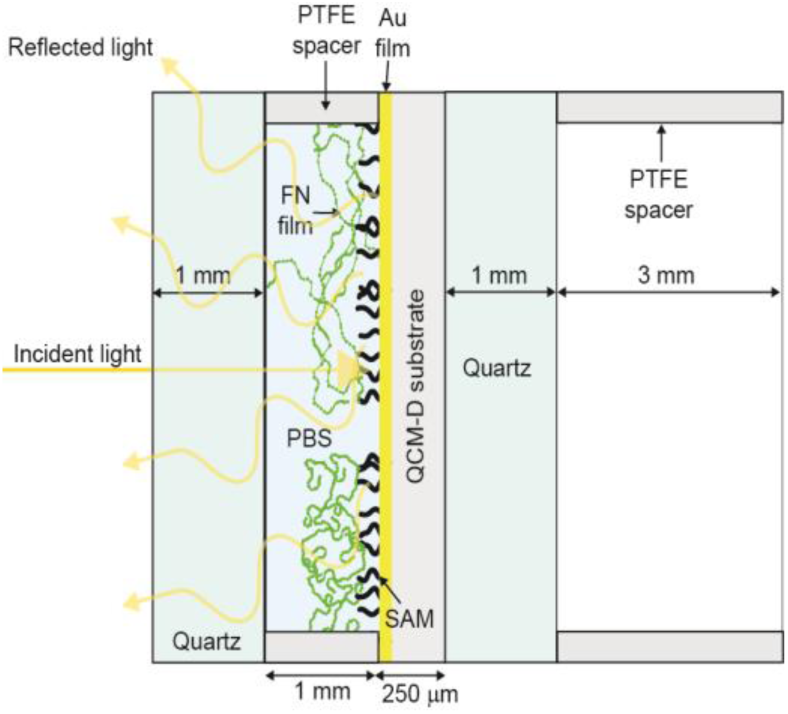
The diffuse reflectance circular dichroism (DRCD) sample holder setup.

As the measured ellipticity is concentration-dependent, conversion to mean residue ellipticity (MRE) was done using the following equation:

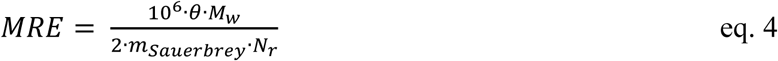

where *θ* is the measured ellipticity in milli degree, *M*_w_ is the protein molecular weight in g/mol (440 kDa for FN),[24] the factor 10^6^ is due to conversion of g to µg, *m*_Sauerbrey_ is the Sauerbrey mass in µg/cm^2^, and *N*_r_ is the number of residues of the adsorbed protein (4956 for FN).[39] In this conversion, we assumed that *m*_Sauerbrey_ is equal to the mass of adsorbed protein per unit area, disregarding the mass of adsorbed solvent. Therefore, the conversion leads to an overestimation. The factor ½ is related to the reflection mode of the measurement setup, which effectively doubles the path length. The entire justification for the conversion, as well as non-converted spectra, can be found in the SI and Figure S5. It should be noted that while conversion of measured ellipticity to MRE based on the path length and sample concentration is a standard practice in liquid-state CD analysis, analogous conversion is rarely performed for solid-state DRCD, albeit precedent does exist.[40,41] The spectra are divided into two regions: far UV (190 nm - 250 nm) and near UV (250 nm - 340 nm), giving indications of protein secondary and tertiary structure, respectively. The far-UV spectra were deconvoluted, and secondary structure analysis was performed using the BeStSel web application (ELTE Eötvös Loránd University, Budapest, Hungary; version fixes 18th March 2024, web server update 13^th^ September 2023).[40,42] The normalized root mean square deviation (NRMSD) was used as a measure of deconvolution uncertainty. Circular dichroism (CD) spectra of bulk FN at a concentration of 50 µg/mL, used to functionalize the gold-coated quartz crystals, are shown in Figure S6 and agree with urea-induced unfolding of FN.[43]

### Atomic Force Microscopy (AFM)

A commercial Cypher VRS AFM (Asylum Research, Santa Barbara, CA, USA) was used for AFM imaging of the FN-functionalized surfaces. It was equipped with PPP-BSI AFM Probes (NanoWorld, Bruker, Santa Barbara, USA), having a nominal spring constant of 0.1 N/m, and was calibrated using the thermal noise method.[44,45] Contact mode measurements, for which these biological probes are optimal due to their low spring constant, were performed in a non-controlled environment at several spots near the center of the surface, with a scan rate between 0.8 and 1 line/sec. No signs of material-dragging due to tip-sample interactions were spotted. Images were processed using the software Gwyddion,[46] version 2.6.7. Raw images were first treated with a second-degree polynomial row alignment, followed by horizontal scar corrections. Cross-sections were taken at specific points of each image, as shown in Figure 5, using Gwyddion to analyze the topographical differences for each condition.

### QCM-D data analysis and statistics

Data analysis was performed using a combination of OriginLab Pro 2017 and Microsoft Excel. Box-and-whisker plots indicate the median and lower and upper quartiles. Each data point (one independent measurement) corresponds to an average of data points collected during the last two minutes of the steady state of the relevant condition using pyQCM-BraTaDio.[36] An independent two-sample Student’s *t*-test (unpaired) was performed as a statistical test to evaluate whether the means of three or more independent measurements of two groups were significantly different from each other (*α* = 0.05), calculated using a Python script written with AI assistance using Scipy API and tkinter.[47,48] If the *p*-value of the test was 0.05 or less, *p* < 0.05, the means were considered significantly different between the two groups. QCM-D_data_statistics.xlsx file contains the summary of the statistical analysis.

## RESULTS AND DISCUSSION

### Self-assembled monolayer length influences FN film formation

It is established that surface chemistry has an impact on the molecular conformation of FN.[33,49–51] To test if the pH-induced FN molecular conformational change induced in solution led to different FN molecular films, SAMs formed on Au-coated quartz crystals were used as substrates and immobilize FN. Au surfaces were functionalized with a short or a long amine-terminated SAM, activated with glutaraldehyde, and exposed to FN solutions, followed by a wash to remove loosely bound FN molecules. For short SAMs, at pH 7 pre-wash, the *m*_Sauerbrey_ of FN films was 500 ± 100 ng/cm^2^, which was close to 40% of the *m*_Sauerbrey_ of FN films at pH 4 pre-wash, 1190 ± 40 ng/cm^2^, Figure 3a. A similar trend was observed for FN films on long-SAMs, for which at pH 7, *m*_Sauerbrey_ of FN films was 880 ± 80 ng/cm^2^, ≈ 30% less than *m*_Sauerbrey_ of FN films at pH 4, 1270 ± 50 ng/cm^2^, Figure 3b. The average *m*_Sauerbrey_ and corresponding standard error of the mean (SEM) of the pre- and post-wash *m*_Sauerbrey_ of adsorbed FN films at pH 7 and 4 are summarized in Table 1, and the corresponding quartiles and median are reported as box and whisker plots in Figure 3.

**Figure 3.**
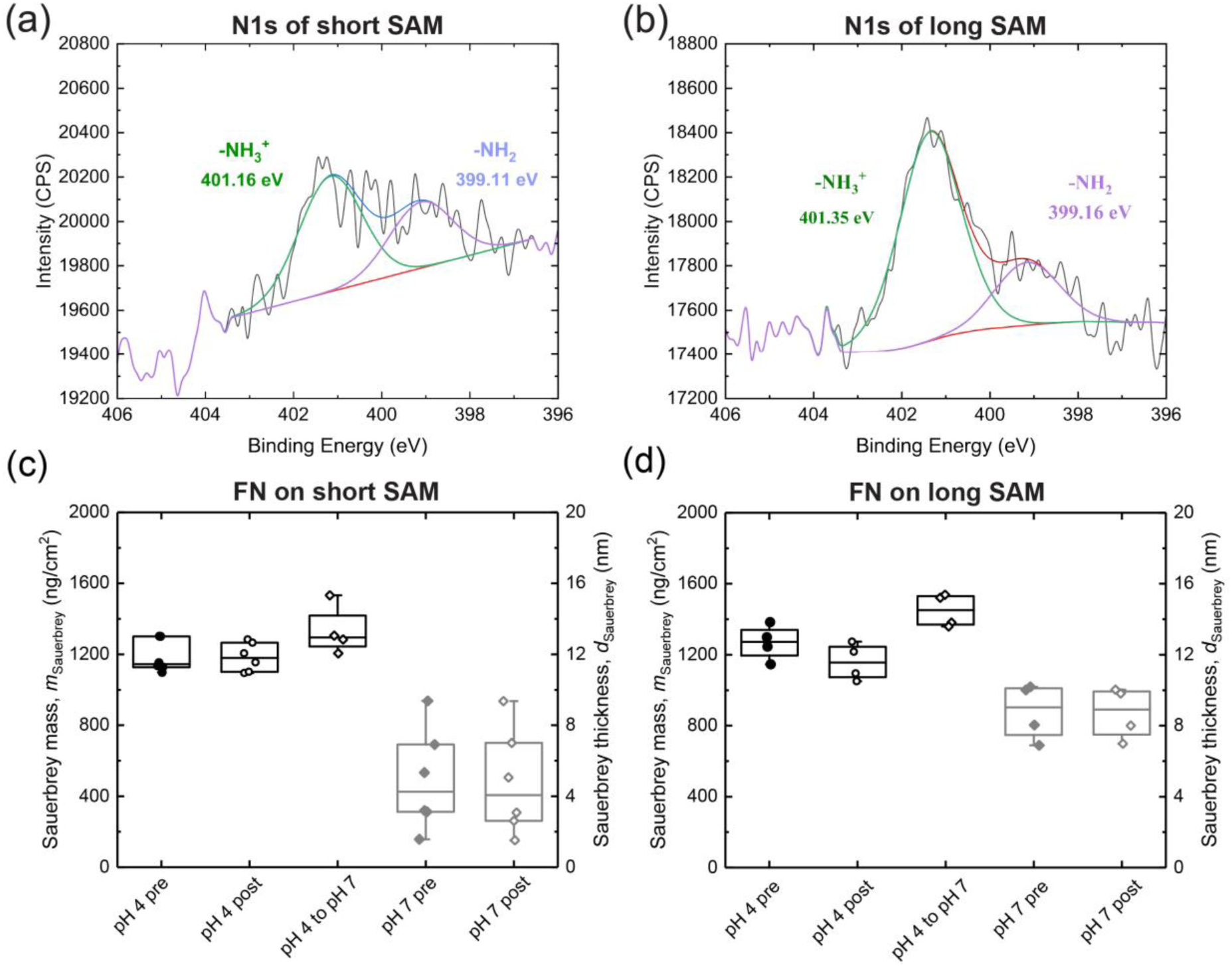
High-resolution XPS spectra of N1s for gold coated QCM-D crystals for (a) short and (b) long SAM, respectively. Long/Short SAM ratio of amine density was found to be around 1.5. Sauerbrey mass *m*Sauerbrey of (c) FN films formed on short SAMs and (d) FN films formed on long SAMs at pH 4, before pH 4 adjusted PBS wash (pH 4 pre), after a pH 4 adjusted PBS wash (pH 4 after), after switching from pH 4 to pH 7 (pH 4 to pH 7), at pH 7 before pH 7 wash, and after a pH 7 PBS wash.

**Table 1.**
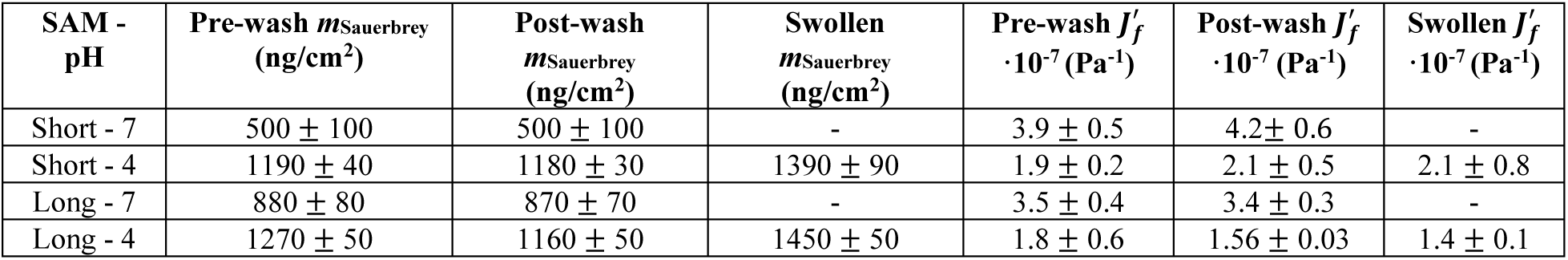
Summary of QCM-D determined FN film properties before (pre) and after (post) PBS washes.

Following the wash with either PBS buffer at pH 7 or pH 4, FN films on short SAMs retained their initial *m*_Sauerbrey_ (pre), being 500 ± 100 ng/cm^2^ for FN on short SAM at pH 7 and 1180 ± 30 ng/cm^2^ for FN on short SAM at pH 4, respectively, corresponding to a *m*_Sauerbrey_ loss of less than 1% for either condition. FN films on long SAMs retained their initial *m*_Sauerbrey_ (pre), being 870 ± 70 ng/cm^2^ for FN on long SAM at pH 7 and 1160 ± 50 ng/cm^2^ for FN on long SAM at pH 4, respectively, corresponding to a *m*_Sauerbrey_ loss of less than 1% for either condition. Switching from pH 4 to pH 7 resulted in swelling of the deposited FN films, with a swollen *m*_Sauerbrey_ of 1390 ± 90 ng/cm^2^ for FN on short SAMs and 1450 ± 50 ng/cm^2^ on long SAMs, respectively. These values correspond to an 18% and 25% increase in *m*_Sauerbrey_ from the PBS 4 post-wash condition for FN films on short and long SAMs, respectively.

Overall, from *m*_Sauerbrey_ or *d*_Sauerbrey_, it is evident that the initial molecular FN conformation impacts molecular FN assembly to form a molecular thin film. “Thicker” films resulted from pre-unfolded FN induced by an acidic pH change. Urea [43] and guanidinium [2] could have been used in previous studies as denaturants; however, these solvents possess densities and viscosities different than PBS,[52,53] inducing undesired baseline shifts in the QCM-D measurements, which were minimal when switching from PBS 4 to PBS 7, as shown in Figure S7. FN binds itself via FN_I_ and FN_III_ repeats near the N- and C-termini,[54] respectively. These domains became more accessible to other FN molecules upon unfolding at acidic pH, leading to increased FN-FN binding and, thus, FN films with more FN molecules. Considering a molecular weight of ⁓500 kDA for one dimeric FN molecule and assuming that the *m*_Sauerbrey_ is due to contributions of the FN molecules alone without water, back-of-the-envelope calculations estimate more than twice the number of FN molecules on films formed from pre-unfolded FN, ⁓14,000 FN molecules per µm^2^ for pH 4 and ⁓6,000 FN molecules per µm^2^ for FN at pH 7 buffer. This is an overestimation, as water molecules contribute to the total sensed *m*_Sauerbrey_ or *d*_Sauerbrey_.[37]

The length of the SAM had an impact on *m*_Sauerbrey_ or *d*_Sauerbrey_, for films formed in PBS 7, but not PBS 4. The Au substrate surface roughness, with an AFM measured root mean square (RMS) surface roughness of ⁓3 nm over a 2 x 2 µm^2^ area, Figure S8a and S8b, was much larger than the short SAM length, estimated to be 0.6 nm (6 carbon-carbon bonds, equivalent to short SAM + glutaraldehyde). It is possible that, on short SAM, partially expanded FN molecules (pH 7) immobilized at the convex summits of the Au surface grains participated in the FN film formation, while partially expanded FN molecules immobilized at the concave valleys of the Au surface grains were less accessible to FN binding domains to other FN molecules in solution, therefore not contributing to the scaffolding of a thick FN film. However, on long SAMs, estimated to be 1.8 nm (15 carbon-carbon bonds, equivalent to long SAM + glutaraldehyde), FN molecules immobilized at the concave valleys of the Au became more accessible to FN binding domains of FN molecules in solution, as longer SAMs smoothened the Au topography and prevented FN molecules from being trapped in surface “crevasses” or valleys. The surface chemistry is known to impact fibronectin assembly and conformation.[55,56] In this case, the difference in amine densities may have played an important role in FN adsorption as well. XPS measurements for long SAMs revealed an amine density ⁓50% higher when compared to the amine density for short SAMs. We suggest that a combination of these two effects led to the measured increase from ⁓6,000 FN molecules per µm^2^ for the short SAMs to ⁓10,000 FN molecules per µm^2^ for the long SAMs.

When switching from PBS 4 to PBS 7, an increase in *m*_Sauerbrey_ or *d*_Sauerbrey_ was observed for FN films on both lengths of SAMs. The swelling could be attributed to conformational changes of FN molecules with sufficient degrees of freedom to accommodate the new spatial molecular arrangement, imposed by changes in intermolecular and intramolecular interactions, leading to water uptake, or increased solvation. FN’s isoelectric point, pI, is predicted to be at a pH of 5.15, [57] with an overall positive charge at pH 4 and a domain-specific charge at neutral pH.[34] The new surface electro-force fields force a more energetically favorable FN inter- and intramolecular interaction and solvation.

### Thin film elastic compliance of FN films tuned by pH

The thin film elastic compliance, *J́*_f,_ was calculated from the slope of ΔΓ vs. − *n* × Δ*f*, as shown in Figure S3b, and described in eq. 2, assuming that η_bulk_ ≈ η_water_.[37] *J*^f́^ of FN films deposited on short SAM at pH 7 (pre) and pH 4 (pre) were (3.9 ± 0.5) ·10^-7^ Pa^-1^ and (1.9 ± 0.2) ·10^-7^ Pa^-1^, respectively, corresponding to a *J́*_f_ difference of ≈ 50%, as shown in Figure 4a. On long SAMs at pH 7 (pre) and pH 4 (pre), *J*^f́^ of FN films were (3.5 ± 0.4) ·10^-7^ Pa^-1^ and (1.8 ± 0.6) ·10^-7^ Pa^-1^, respectively, corresponding to a *J́*_f_ difference of ≈ 50% as well, shown in Figure 4b.

**Figure 4.**
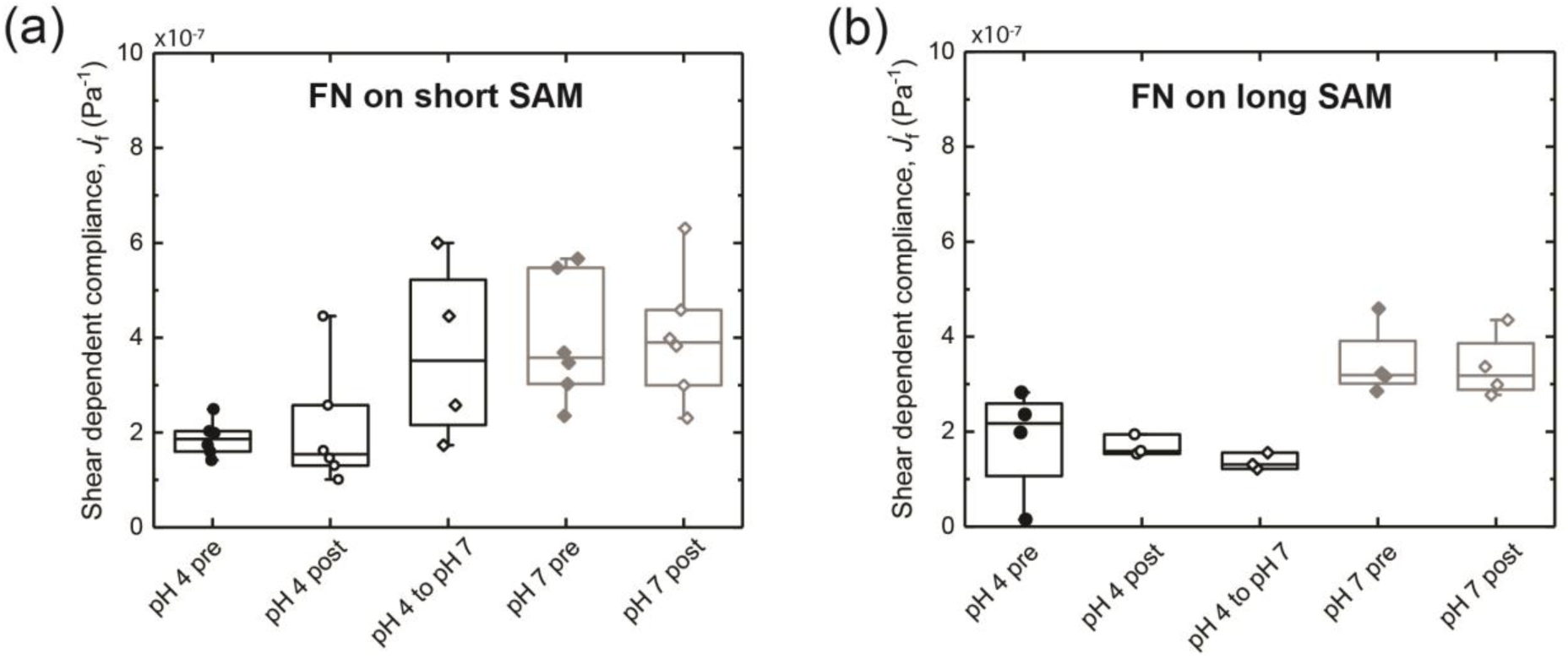
Shear dependent elastic compliance *J*^′^_f_ of (a) FN films formed on short SAMs and (b) FN films formed on long SAMs at pH 4, before pH 4 adjusted PBS wash (pH 4 pre), after a pH 4 adjusted PBS wash (pH 4 after), after switching from pH 4 to pH 7 (pH 4 to pH 7), at pH 7 before pH 7 wash, and after a pH 7 PBS wash.

After the pH 7 and pH 4 washes (post), FN films on short SAMs retained *J*^f́^ values similar to the pre-wash elastic compliances, being (4.2 ± 0.6) ·10^-7^ Pa^-1^ and (2.1 ± 0.5) ·10^-7^ Pa^-1^, for pH 7 and pH 4, respectively, corresponding to a *J́*_f_ difference of ≈ 5% and ≈ 10% relative to the pre-wash FN films, Figure 4a. FN films on long SAMs also retained their initial *J́*_f_, being (3.4 ± 0.3) ·10^-7^ Pa^-1^ and (1.56 ± 0.03) ·10^-7^ Pa^-1^ for pH 7 and pH 4, respectively, Figure 4b. Here, the *J́*_f_ differences from the pre-wash FN films were ≈ 3% and ≈ 10%. Table 1 summarizes the average *J́*_f_ and corresponding SEM of the pre- and post-wash *J́*_f_ of adsorbed FN films at pH 7 and 4. The corresponding quartiles and medians are reported as box and whisker plots in Figure 4.

We discuss the results in terms of *J́*_f_, as it contains contributions of both collected signals, Δ*f*_n_ and Δ*D*_n_, for all available overtones. That is because the shift of dissipation of a QCM-D substrate when operating in liquid caused by a thin film is proportional to the elastic compliance of the film.[37,58] *J́*_f_ showed similar trends for FN films on short and long SAMs—less compliant elastic films formed at pH 4 and more compliant ones formed at pH 7. The results can be interpreted as follows: changes in tertiary and secondary structure, as measured by DRCD and discussed below, lead to less compliant FN films, probably due to the loss of the *β*-barrel and *β*-sheet contents at pH 4, while more tertiary and secondary order, as is the case for partially extended FN molecules at pH 7, assembled into more compliant structures with more *β*-barrel and *β*-sheet content. Intriguingly, differences between short and long SAMs became evident in the transition from pH 4 to pH 7. For FN films on short SAMs, *J́*_f_ increased to similar values to those formed at pH 7. This change in *J́*_f_ suggests molecular conformational flexibility, allowing for adjustments to inter- and intramolecular interactions imposed by solvation and electrostatics. That was not true for FN films on long SAMS, which retained their pH 4 imposed *J́*_f_. It appears that on long SAMS, the initial FN film molecular conformation was retained or frozen, and FN molecules lacked the conformational flexibility to adjust for solvation forces and electrostatics.

To test the swelling and change in elastic compliance of the FN films observed during the PBS 4 to PBS 7 transition, control experiments using bare Au substrates were conducted to quantify potential contributions in baseline shift due to potential viscosity and density changes. The buffer-induced change in frequency signal was under 2.5 Hz, Figure S7, significantly lower than the swelling response, −45 Hz, concluding that the signal changes resulted from FN molecular conformational changes.

AFM imaging is reported next to obtain visual information on FN film morphologies deposited on short and long SAMs with PBS at pH 7 and PBS at pH 4.

### AFM reveals morphological differences of surface-bound FN

AFM imaging of FN films was used to visualize morphological similarities and differences induced by the pH and SAM length. AFM friction imaging of FN films revealed differences in morphology due to both explored parameters, deposition pH and SAM length, as shown in Figure 5. Lateral force (with millivolt as unit) revealed finer morphological details than the height channel, Figure S8. For all conditions, AFM reveals that FN films completely covered the surfaces – in agreement with results shown in Figure S4, which indicates that full saturation has been reached at the working concentration. FN film morphologies on short SAMs, independent of pH, Figure 5a and 5b, possessed higher peak-to-valley variations than FN film morphologies on long SAMs, independent of pH, Figure 5c and 5d. Additionally, peak-to-valley variations were larger for FN film morphologies formed with PBS 7 than for FN films formed with PBS 4, irrespective of SAM length.

**Figure 5.**
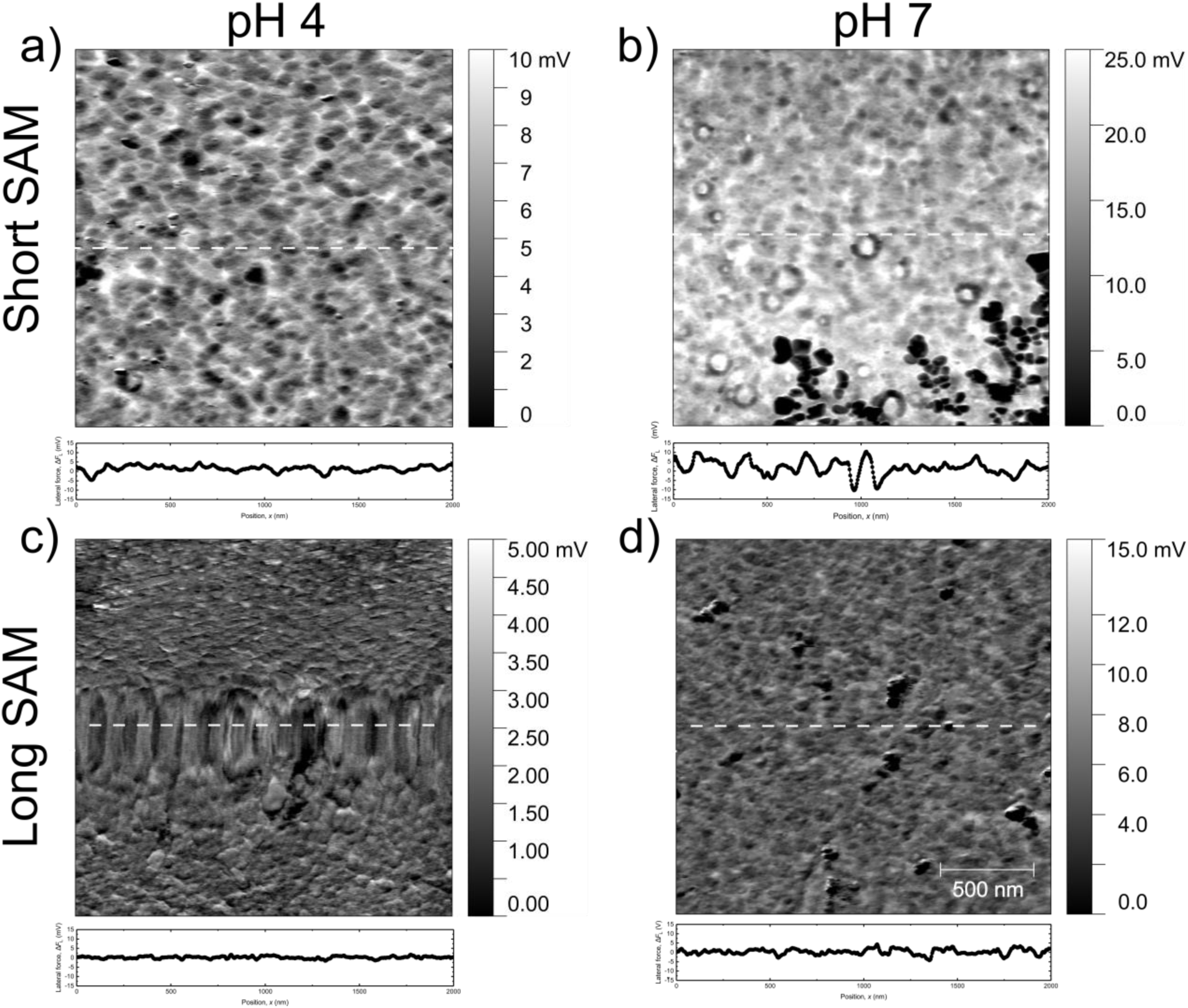
AFM friction maps of FN films at four different conditions: (a) Short SAM pH 4, (b) Short SAM pH 7, (c) Long SAM pH 4, (d) Long SAM pH 7. Cross-sections of representative spots for each case are shown, which can be understood as differences in the tip-sample interaction/friction. The 500 nm scalebar shown applies to all panels.

In short SAMs, the film morphology was dominated by the gold roughness for pH 4 and 7, - Figures 5a and 5b and Figure S8a and S8b for comparison with a bare gold surface. This result is not surprising, as the short SAMs consisted of a 2-carbon chain, with an expected height close to one-fifth of the bare gold surface root mean squared (RMS) roughness, which was measured to be approximately 3 nm for a 2 x 2 µm² area, as shown in Figure S8a. Long SAMs, however, appeared to be long enough to prevent the underlying gold granular topography from dominating the FN film morphology. This effect is complementary to the difference in amine density shown by XPS, which is likely to dominate FN adsorption. A unique feature observed for pH 4 conditions, particularly on long SAMs, was the presence of two domains: one relatively smooth with small globular domains and a second with elongated “grains”. These elongated grains were not observed in PBS 7 samples on either short or long SAMs. They mostly appeared in the slow-scan axis of the AFM, *i.e.*, perpendicular to the “dragging” direction of the tip, from where we don’t consider them as measuring artifacts. Differences in the cross sections shown in Figure 5 must be considered as heterogeneous tip-sample interactions but not be taken as a roughness measurement, as they are not topographical (height) maps.

AFM images of films after adsorption of synovial fluid molecules were not collected, as the focus was on characterizing FN absorbed films. DRCD spectroscopy was conducted to obtain quantitative information on the secondary structure of the FN films deposited on short and long SAMs with PBS 7 or PBS 4, as described next.

### Beta sheet content changes are preserved in surface-bound FN

DRCD analysis in the far UV range (ca. 190-250 nm) gives insights into protein secondary structure.[59] Here, typical peaks associated with *β*-sheets between 200-210 nm are observed, as shown in Figure 6a and 6c.[60] This was expected due to the strong tendency of fibronectin to form *β*-sheet-rich structures in its native form.[61] The deconvolution results (Table 2) indicate that the adsorption pH and SAM type have some effect on the secondary structure of the adsorbed proteins, with pH 7 favoring *β*-sheets on both short and long SAMs. Interestingly, deconvolution of FN adsorbed on a long SAM at pH 4 showed that *α*-helices dominate over *β*-sheets. It can be expected that when adsorbed on long SAM, the surface effect influence on FN structure is minimized, suggesting that at short SAM, 0.6 nm long, we observed a decrease in *α*-helices content due to van der Waals forces acting on the protein chain on top of the pH-induced structure changes. pH is known to influence the conformation of free FN in suspension.[34] Based on DRCD, and consistent with QCM-D, it appears that on long SAMs, FN’s molecular conformation imposed by bulk pH was preserved in immobilized FN molecules. It should also be noted that a significantly stronger MRE signal was observed at pH 7 compared to pH 4; we attribute this to the unfolding of FN at pH 4, as discussed below.

**Figure 6.**
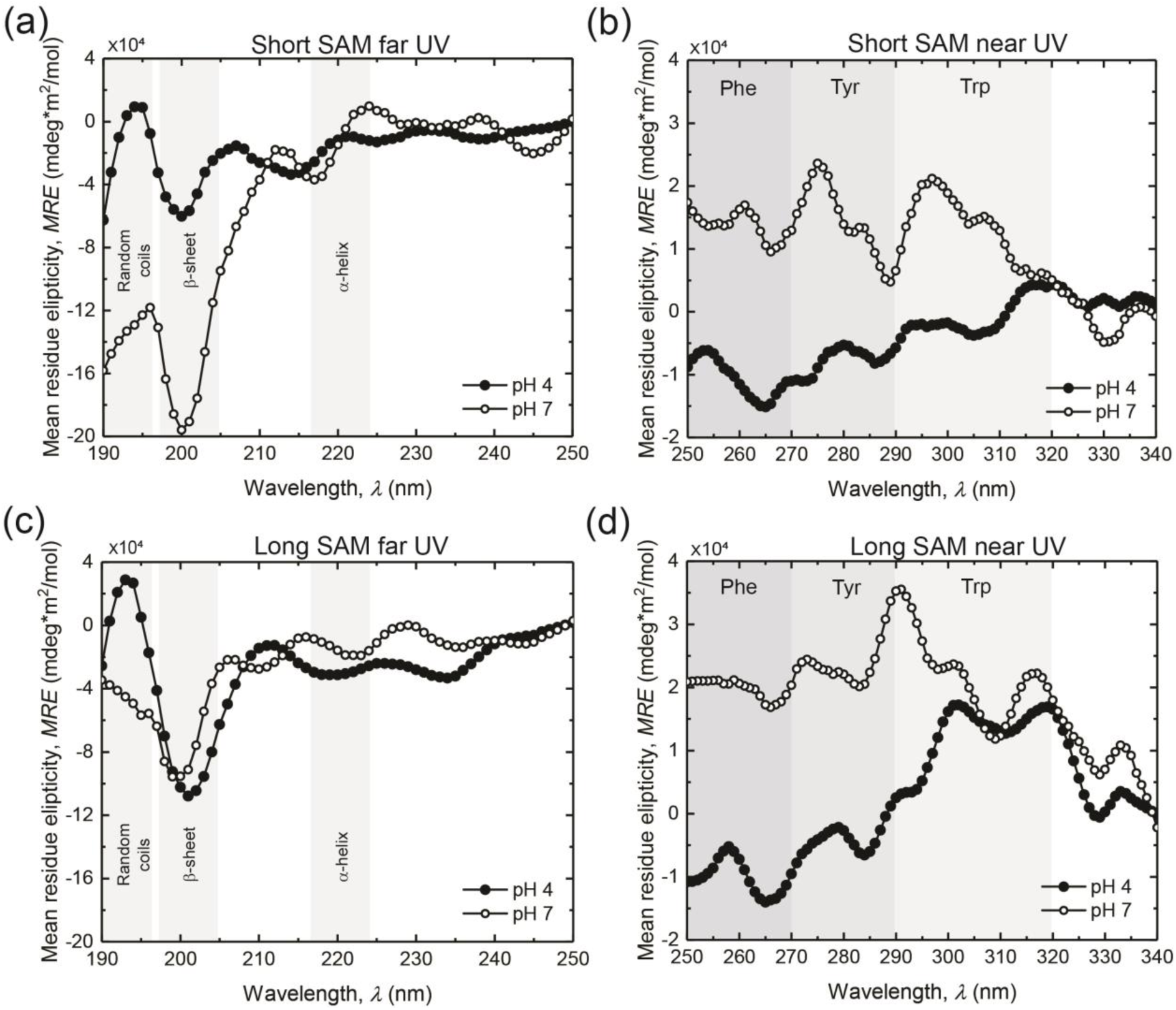
(a) Far UV DRCD spectra of FN adsorbed onto a short SAM at pH 4 and pH 7, reported as mean residue ellipticity, MRE, (b) far UV DRCD spectra of FN adsorbed onto a long SAM at pH 4 and pH 7 in MRE, (c) near UV DRCD of FN adsorbed onto short SAM at pH 4 and pH 7 in MRE, and (d) near UV DRCD of FN adsorbed onto a long SAM at pH 4 and pH 7 in MRE.

**Table 2.** Secondary structure content as quantified via DRCD.

| SAM length | pH | α-helices | β-sheets | Turns | Other | NRMSD |
| --- | --- | --- | --- | --- | --- | --- |
| Short | 4 | 8.9 | 35.9 | 0.5 | 54.6 | 0.16 |
| Short | 7 | 13.3 | 77.5 | 6.2 | 2.9 | 0.3 |
| Long | 4 | 41.4 | 12.0 | 0.0 | 46.6 | 0.17 |
| Long | 7 | 12.2 | 58.4 | 9.4 | 20.0 | 0.14 |

DRCD analysis in the near-UV range (250-340 nm) can provide insights into protein tertiary structure based on the interaction of aromatic residues (*e.g.*, tyrosine and tryptophan), as shown in Figures 6b and 6d. Near-UV DRCD can detect conformational changes within protein chains, as well as the arrangement of disulfide bonds, hydrogen bonding, and the mobility of each amino acid residue.[60,62] Here, FN layers exhibited a broad positive peak, indicating a certain level of tertiary structure ordering.[60,63] FN exhibited two peaks, clearly visible between 270-290 nm and 290-320 nm, as shown in Figures 6b and 6d, corresponding to the absorbance of both tyrosine and tryptophan. Notably, there was a shift in the peak positions between the long and short SAM, with peaks tending to occur at higher wavelengths for FN adsorbed on the long SAM. This suggests a difference in either hydrogen bonding or conformational flexibility for aromatic segments on the different substrates.[60] However, as hydrogen bonding is expected to be similar at the same pH on both long and short SAM samples, we attribute this primarily to a change in FN conformational flexibility. These results confirm the observations from QCM-D, where an evident change in *J́*_f_ on short SAMs suggests molecular conformational flexibility, allowing for adjustments to inter- and intramolecular interactions imposed by solvation and electrostatics. That was not true for FN films on long SAMs, which retained their pH 4 imposed *J́*_f_.

Similar to secondary structure, the tertiary structure of FN seems to be affected by the length of SAM. On short SAM, we observed a prominent tyrosine double peak appearing at pH 7, between 275-290 nm alongside a clear tryptophan peak around 300 nm.[60] Both peaks are, however, largely suppressed at pH 4, suggesting a significant shift in the conformation of the protein chains with respect to pH. In particular, this indicates that at pH 4, protein mobility is significantly increased, as tertiary organization is disrupted by the random motion of the protein chain segments. This suggests that at pH 4, FN exists largely in a disorganized tertiary structure when adsorbed on a short SAM. For long SAM, however, the tryptophan and tyrosine peaks remain highly prominent in both pH 4 and 7. There is a notable peak shift in tyrosine signal with the change of pH, which suggests a shift in protein tertiary structure, although there is some ordering present at both pH. Furthermore, the phenol peak is more prominent at pH 4 than at pH 7, indicating another structural shift induced by pH. As discussed earlier, while the changes on short SAM can be affected by surface interactions with the gold substrate, these effects should be negligible for long SAM, and the changes observed in DRCD are thus reflective of the behavior of the protein in an adsorbed thin layer.

To assess FN’s role in scaffolding a lubricating and wear-protecting film and FN’s conformation impact in scaffolding a lubricating and wear-protecting film, FN films on short and long SAMs deposited at pH 4 and pH 7 were exposed to dSF in a QCM-D fluid cell and its mass adsorption monitored, as discussed below.

### FN film conformation controls dilute synovial fluid (dSF) adsorption

After flowing BSA to block non-specific binding, the amount of diluted synovial fluid (dSF) adsorbed on FN films was estimated via QCM-D. On short SAMs, dSF *m*_Sauerbrey_ was 210 ± 60 ng/cm^2^ on pH 7 deposited FN films. However, on FN films deposited on short SAMs at pH 4, dSF *m*_Sauerbrey_ was 160 ± 50 ng/cm^2^, ≈ 25% less than on FN films deposited at pH 7. After the PBS wash, always done with PBS 7, *m*_Sauerbrey_ of dSF on pH 7 and pH 4 deposited FN films were 160 ± 50 ng/cm^2^ and 80 ± 30 ng/cm^2^, respectively. These values correspond to a *m*_Sauerbrey_ decrease of 24% and 50% relative to their corresponding prewash *m*_Sauerbrey_, respectively.

On long SAMs, dSF *m*_Sauerbrey_ was 90 ± 20 ng/cm^2^ on pH 7 deposited FN films. On FN films deposited on long SAMs at pH 4, dSF *m*_Sauerbrey_ was 90 ± 10 ng/cm^2^, about the same as that on pH 7 deposited FN films. After the PBS wash, *m*_Sauerbrey_ of dSF on pH 7 and pH 4 deposited FN films were 60 ± 10 ng/cm^2^ and −2 ± 20 ng/cm^2^ (or no adsorption), respectively. These values correspond to a *m*_Sauerbrey_ decrease of ≈ 30% and ≈ 100% relative to their corresponding prewash *m*_Sauerbrey_, respectively. Table 3 summarizes the average *m*_Sauerbrey_ and corresponding SEM of the pre- and post-wash dSF adsorbed onto FN films formed on short and long SAMS deposited at pH 7 and 4. The corresponding quartiles and medians are reported as box and whisker plots in Figure 7.

**Figure 7.**
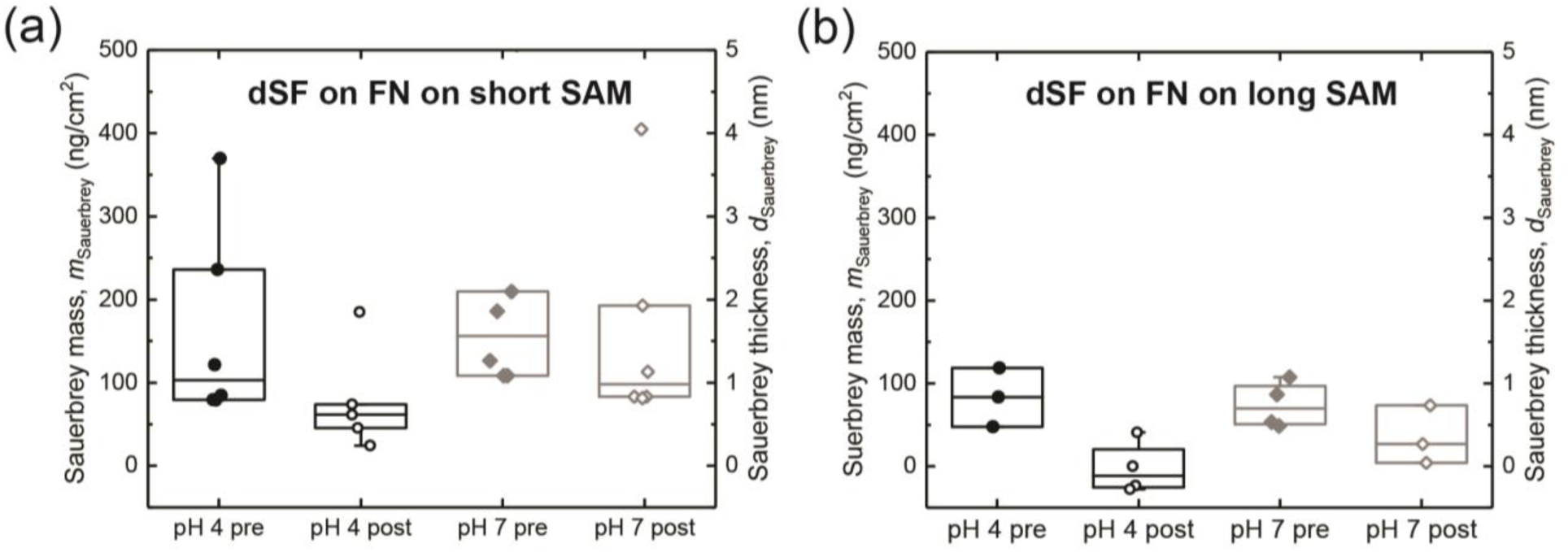
Sauerbrey mass *m*Sauerbrey of (a) dSF on FN films formed on short SAMs and (b) dSF on FN films formed on long SAMs, before (pre) and after (post) a PBS 7 wash.

**Table 3.** Summary of QCM-D determined dSF film properties before (pre) and after (post) PBS washes.

| FN on SAM - pH | Pre-wash dSF $m_{\text{Sauerbrey}}$ (ng/cm<sup>2</sup>) | Post-wash dSF $m_{\text{Sauerbrey}}$ (ng/cm<sup>2</sup>) | Pre-wash dSF $J_f'$ $\cdot 10^{-7} (\text{Pa}^{-1})$ | Post-wash dSF $J_f'$ $\cdot 10^{-7} (\text{Pa}^{-1})$ |
| --- | --- | --- | --- | --- |
| Short - 7 | $210 \pm 60$ | $160 \pm 50$ | $8 \pm 4$ | $6 \pm 2$ |
| Short - 4 | $160 \pm 50$ | $80 \pm 30$ | $30 \pm 10$ | $12 \pm 4$ |
| Long - 7 | $90 \pm 20$ | $60 \pm 20$ | $16 \pm 2$ | $40 \pm 10$ |
| Long - 4 | $90 \pm 10$ | No adsorption | $60 \pm 30$ | No adsorption |

Control experiments consisted of quantifying *m*_Sauerbrey_ of dSF on bare Au surfaces, Au surfaces with short and long SAMs, and Au surfaces with short and long SAMs and a BSA blocking step, to gain insights into the role of FN films in adsorbing and retaining dSF, Figure S9 and S10. In all these control conditions, *m*_Sauerbrey_ of dSF was significantly higher than *m*_Sauerbrey_ of dSF on FN films, indicating that the dSF on FN films is due to contributions of solely FN films and not underlying bare gold or SAM surfaces.

The amount of dSF adsorbed to and retained by the FN films is significantly less than what has been reported to negatively charged mica surfaces, [64] by almost 1000 ng/cm^2^. The observation suggests that FN films saturate at much lower SF concentrations and that component adsorption to FN was either different, oriented differently, or a combination of both relative to when compared to mica, a commonly used model substrate. The primary proteinaceous constituent of SF is serum albumin,[65] which was used as a non-specific blocker in this study. On clean, bare Au surfaces (no SAM and no FN), BSA formed a film of *m*_Sauerbrey_ = 1000 ± 100 ng/cm^2^ and *m*_Sauerbrey_ = 1000 ± 100 ng/cm^2^ a, pre- and post-wash with PBS 7, respectively. Similar adsorption magnitudes of BSA onto Au have been reported in other studies.[66,67] However, on FN films, the amount of BSA adsorbed and retained decreased considerably to 120 ± 10 ng/cm^2^ and 380 ± 50 ng/cm^2^, for FN films formed on short and long SAM post-wash formed at a pH of 4. Conversely, the amount of BSA adsorbed and retained decreased to 140 ± 20 ng/cm^2^ and 90 ± 20 ng/cm^2^, for FN films formed on short and long SAM post-wash formed at a pH of 7. These results are summarized in Figure S10 and Table S1. Combined, it is evident that FN films mediate the adsorption of dSF, including BSA.

To account for the differences in *m*_Sauerbrey_ of the various FN films and its contributions in retaining dSF, for example due to the number of molecules participating in dSF adsorption and retention, the *m*_Sauerbrey_ of dSF (pre- and post-wash) was normalized by the *m*_Sauerbrey_ of FN quantified before the dSF step. For FN films deposited with PBS 4, the *m*_Sauerbrey_ used for normalization was measured after the switch from pH 4 to pH 7. Equivalently, for FN films deposited with PBS 7, the *m*_Sauerbrey_ used for normalization was measured for FN films after PBS 7 wash, Figure 8. Based on this normalization, less dSF adsorbed (pre) and was retained (post) on FN films formed on long SAMs, independent of pH, than on FN films formed on short SAMs. For the PBS 4 condition, dSF adsorption on FN on long SAMs was 54% less than on FN on short SAMs pre-wash. On average, no adsorption was measured on FN on long SAMs post-wash. For the PBS 7 conditions, dSF adsorption on FN on long SAMs was 87% and 93% less than on FN on short SAMs, pre- and post-wash, respectively. Normalization also revealed that less dSF adsorbed to FN films formed with PBS 4 on short and long SAMs, when compared to their PBS 7 counterparts. On FN films deposited on short SAMs at pH 4, dSF adsorption was 80% and 90% less than on FN deposited at pH 7, pre- and post-wash, respectively. On FN films deposited on long SAMs, dSF adsorption with PBS 4 was 25% less than with PBS 7, prewash. No dSF remained adsorbed after the wash on FN films on long SAMs formed with PBS 4.

**Figure 8.**
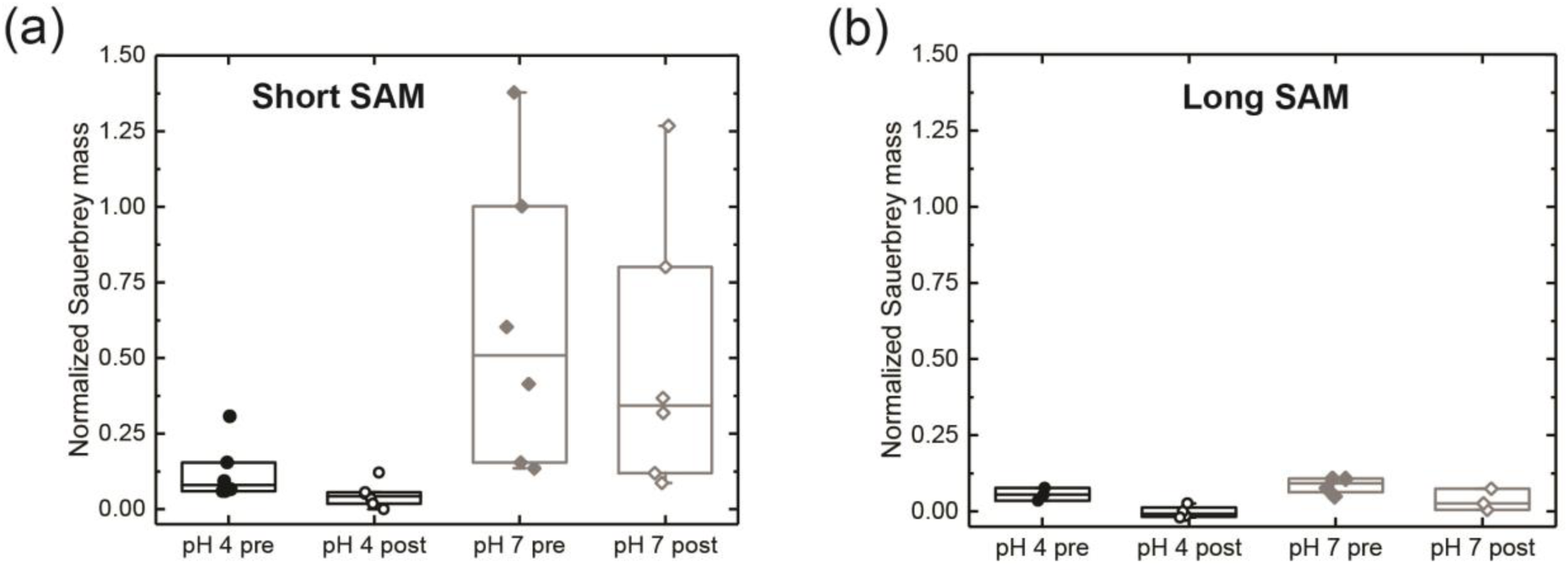
Normalized *m*Sauerbrey of dSF over *m*Sauerbrey of FN (a) for short SAMs conditions and (b) long SAMs conditions.

Overall, as supported by DRCD spectroscopy, FN films formed at a pH of 7, independent of SAM length, with more β-sheet content and tertiary structure, retain more dSF than FN films formed at pH of 4. This is the first report showing evidence that FN’s conformation at the surface of articular cartilage could play a role in the assembly process of a wear-protecting and lubricious film, and that FN could be an overlooked important molecular regulator of articular cartilage surface mechanical homeostasis. It becomes evident that identifying the component that binds the strongest and with the highest affinity to the various FN conformations is crucial to assist in the design of molecular therapies to recover function in pathophysiology, such as in osteoarthritis. Lubricin (or also PRG4) has been identified as a crucial boundary lubricant.[23,68–72]. However, binary,[26,29,73] ternary, or higher-order synergistic interactions [64,74] are most likely behind the remarkable tribological properties of synovial joints. To identify them, higher throughput assays will have to be developed and implemented. Furthermore, how other macromolecular constituents of the articular cartilage ECM interact with FN in unison with SF components adds to the complexity of isolating crucial synergies. Therefore, it will be essential to investigate how SF adsorption changes to FN in the presence of collagen II, as FN/collagen II interactions can drastically alter the SF adlayers, as known for other SF and articular cartilage ECM macromolecular constituents, such as lubricin-hyaluronan, hyaluronan-aggrecan complexes, or hyaluronan-phospholipids. [31]

### Thin film elastic compliance of dSF adsorbed on FN films is affected by loss of dSF

Similarly, the determination of the thin film elastic compliance of dSF films formed on FN was estimated using the previously described approach, with results summarized in Figure 9, assuming that η_*dSF*_ ≈ η_water_. *J*^f́^ of dSF films deposited on FN on short SAM at pH 7 (pre) and pH 4 (pre) were (8 ± 4)·10^-7^ Pa^-1^ and (30 ± 10)·10^-7^ Pa^-1^, respectively, corresponding to a 4 times increase in *J́*_f_, as shown in Figure 9a. On FN on long SAMs at pH 7 (pre) and pH 4 (pre), *J*^f́^ of dSF films were (16 ± 1)·10^-7^ Pa^-1^ and (60 ± 30)·10^-7^ Pa^-1^, respectively, corresponding to a similar increase in *J́*_f_, as shown in Figure 9b.

**Figure 9.**
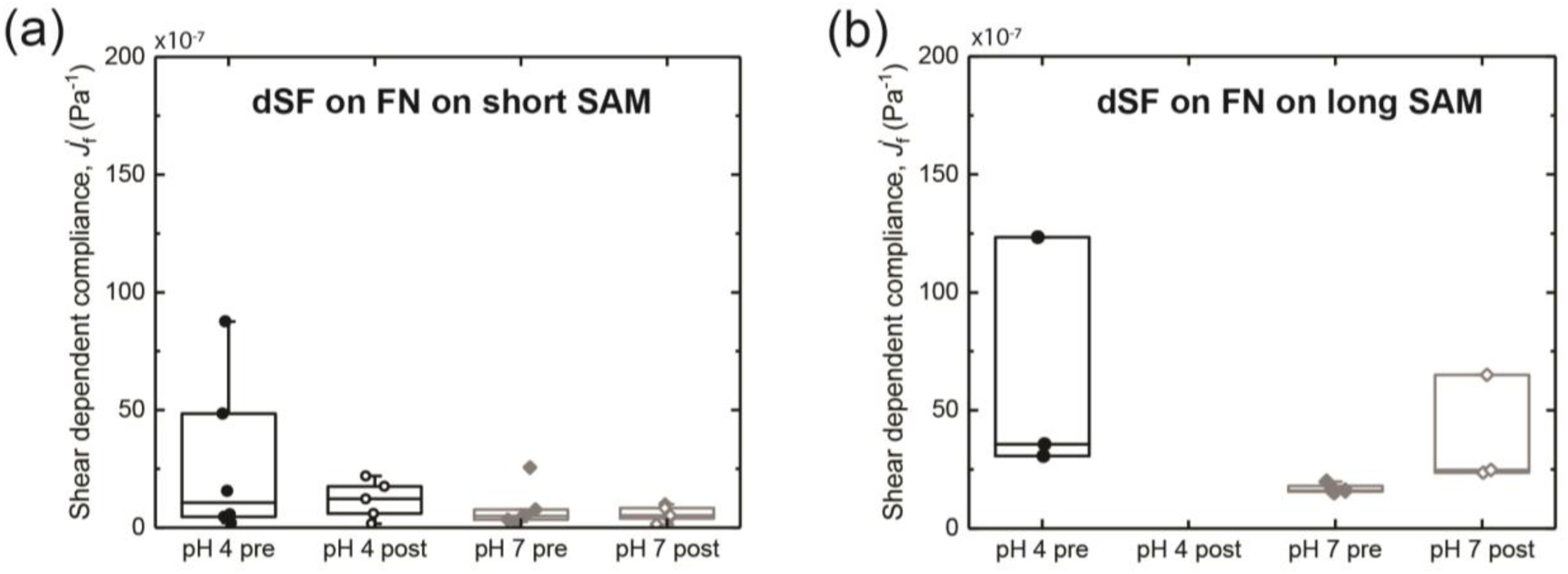
Shear-dependent elastic compliance *J*^′^_f_ of (a) dSF films formed on FN formed on short SAMs and (b) dSF films formed on FN formed on long SAMs. Condition pH 4 post has no shear-dependent compliance, as the adsorption average was zero.

After the pH 7 and pH 4 wash (post-), *J*^f́^ of dSF films on FN films on short SAMs decreased relative to the pre-wash elastic compliances, being (6 ± 2) ·10^-7^ Pa^-1^ and (12 ± 4) ·10^-7^ Pa^-1^, for pH 7 and pH 4, respectively, corresponding to a *J́*_f_ decrease of ≈ 25% and ≈ 60% relative to the pre-wash dSF films, Figure 9a. However, dSF films on FN films on long SAMs had a dramatic change relative to their initial *J́*_f_, being (40 ± 10)·10^-7^ Pa^-1^ and (−80 ± 50)·10^-7^ Pa^-1^ for pH 7 and pH 4, respectively, Figure 9b. Here, the *J́*_f_ differences from the pre-wash FN films were an increase of ≈ 240% and a loss of the films, and thus a negative elastic compliance was obtained. Table 3 summarizes the average *J́*_f_ and corresponding SEM of the pre- and post-wash *J́*_f_ of adsorbed dSF onto FN films formed on short and long SAMs at pH 7 and 4. The corresponding quartiles and medians are reported as box and whisker plots in Figure 9.

Overall, the dSF films adsorbed on FN deposited at pH 4 are more rigid and form a thinner adlayer than those on films deposited at pH 7. The observations are more apparent for short SAM functionalized surfaces. The loss of film results in negative values of thin shear compliance, which is not observed for short SAM functionalized surfaces. dSF films formed on FN films with less tertiary structure and less *β*-sheet content were the most compliant, despite not being the FN conditions that adsorbed the highest dSF. This finding could imply that it is not the amount of components adsorbed to the surface of articular cartilage but rather the compliance that dictates the tribological performance. To elucidate, it will be necessary to conduct nanotribological studies, which are beyond the scope of this study.

## CONCLUSIONS

Results reported in this study support the relevance of understanding FN conformation at the surface of articular cartilage. Findings indicate that unfolded FN (pH 4) molecules formed thicker films with (modest) less *β*-sheet content and smoother morphologies than films formed from extended FN (pH 7) molecules. However, FN films formed from extended FN retained more SF than films formed from unfolded FN molecules. Overall, this study presents a model to investigate FN interacting with SF and assisting in elucidating FN’s role in scaffolding a lubricating and wear-protecting film, demonstrating that FN conformation is involved in mediating chemo-mechanical properties of the articular cartilage ECM and SF interface. This is the first report showing evidence that FN’s conformation at the surface of articular cartilage could play a role in the assembly process of a wear-protecting and lubricious film, and that FN could be an overlooked crucial molecular regulator of articular cartilage surface mechanical homeostasis. This knowledge will allow a better understanding of the molecular regulation of the articular cartilage-SF interface homeostasis, and identify molecular interactions and synergies of articular cartilage ECM and SF to reveal the complexity of synovial joints.

## ACKNOWLEDGMENTS

R.C.A.E., D.R.J.P., and U.H. acknowledge funding from the National Science Foundation (NSF) CAREER Award through NSF-CMMI-2239665 awarded to R.C.A.E.. R.C.A.E., S.T.A., and K.C. acknowledge NSF LEAP HI Award funding through NSF-CMMI-2245367. K.L. and R.C.A.E. acknowledge funding from NSF REU SITES: Interdisciplinary Biological Engineering and Science Training (I-BEST) through NSF-DBI-2349757. R.C.A.E. acknowledges funding from NSF-CREST: Center for Cellular and Biomolecular Machines through the support of NSF-HRD-1547848. K.D. acknowledges funding from the Natural Sciences and Engineering Research Council of Canada (NSERC) Discovery Grants Program (RGPIN-2023-03607). Thanks to Prof. Mehmet Baykara, Gokay Adabasi, and Joshua R. Evans from the Department of Mechanical Engineering at the University of California, Merced, for allowing and coordinating the use of AFM. We thank the Imaging and Microscopy Facility at the University of California, Merced for XPS access, and Dr. Stavros G. Karakalos for enriching discussions on XPS data collection and interpretation.

## SUPPLEMENTARY INFORMATION

Figure S1. Schematics of chemical immobilization of FN to amine-functionalized surfaces with glutaraldehyde.

Figure S2. XPS spectra of the control gold substrate without SAMs.

Figure S3. Graphical determination of Sauerbrey mass, *m*_Sauerbrey_, and thin film compliance, *J*_f_′.

Figure S4. FN film formation as a function of FN bulk concentration and corresponding thin film compliances.

Figure S5. Non-converted DRCD spectra.

Figure S6. Circular dichroism spectra of fibronectin in solution (bulk) at pH 4 and pH 7 Figure S7. QCM-D baseline shifts due to pH change.

Figure S8. AFM images of control surfaces and height images of Figure 5. Figure S9. Sauerbrey mass, *m*_Sauerbrey_, of dSF onto various control surfaces. Figure S10. Sauerbrey mass, *m*_Sauerbrey_, of BSA onto various control surfaces.

Table S1. Values of Sauerbrey mass, *m*_Sauerbrey_, of BSA onto various control surfaces shown in Figure S7.

## REFERENCES

[1] R. Pankov, Fibronectin at a glance, J. Cell Sci. 115 (2002) 3861–3863.

[2] M.L. Smith, D. Gourdon, W.C. Little, K.E. Kubow, R.A. Eguiluz, S. Luna-Morris, V. Vogel, Force-induced unfolding of fibronectin in the extracellular matrix of living cells, PLoS Biol. 5 (2007) e268.

[3] J. Patten, K. Wang, Fibronectin in development and wound healing, Adv. Drug Deliv. Rev. 170 (2021) 353–368.

[4] J. Patten, P. Halligan, G. Bashiri, M. Kegel, J.D. Bonadio, K. Wang, EDA fibronectin microarchitecture and YAP translocation during wound closure, ACS Biomater. Sci. Eng. (2025). 10.1021/acsbiomaterials.4c02019.

[5] A. Krammer, H. Lu, B. Isralewitz, K. Schulten, V. Vogel, Forced unfolding of the fibronectin type III module reveals a tensile molecular recognition switch, Proc. Natl. Acad. Sci. U. S. A. 96 (1999) 1351–1356.

[6] A.J. Zollinger, M.L. Smith, Fibronectin, the extracellular glue, Matrix Biol. 60–61 (2017) 27–37.

[7] K. Wang, R.C. Andresen Eguiluz, F. Wu, B.R. Seo, C. Fischbach, D. Gourdon, Stiffening and unfolding of early deposited-fibronectin increase proangiogenic factor secretion by breast cancer-associated stromal cells, Biomaterials 54 (2015) 63–71.

[8] E. Zamir, B.Z. Katz, S. Aota, K.M. Yamada, B. Geiger, Z. Kam, Molecular diversity of cell-matrix adhesions, J. Cell Sci. 112 (pt11) (1999) 1655–1669.

[9] E. Rouslahti, Fibronectin and its receptors, Annu. Rev. Biochem. 57 (1988) 375–413.

[10] E. Klotzsch, M.L. Smith, K.E. Kubow, S. Muntwyler, W.C. Little, F. Beyeler, D. Gourdon, B.J. Nelson, V. Vogel, Fibronectin forms the most extensible biological fibers displaying switchable force-exposed cryptic binding sites, Proc. Natl. Acad. Sci. U. S. A. 106 (2009) 18267–18272.

[11] E.M. Chandler, B.R. Seo, J.P. Califano, R.C. Andresen Eguiluz, J.S. Lee, C.J. Yoon, D.T. Tims, J.X. Wang, L. Cheng, S. Mohanan, M.R. Buckley, I. Cohen, A.Y. Nikitin, R.M. Williams, D. Gourdon, C. a. Reinhart-King, C. Fischbach, Implanted adipose progenitor cells as physicochemical regulators of breast cancer, Proc. Natl. Acad. Sci. U. S. A. 109 (2012) 9786–9791.

[12] R. Teshima, M. Ono, Y. Yamashita, H. Hirakawa, K. Nawata, Y. Morio, Immunohistochemical collagen analysis of the most superficial layer in adult articular cartilage, J. Orthop. Sci. 9 (2004) 270–273.

[13] R. Crockett, A. Grubelnik, S. Roos, C. Dora, W. Born, H. Troxler, Biochemical composition of the superficial layer of articular cartilage, J. Biomed. Mater. Res. A 82 (2007) 958–964.

[14] M. Watanabe, C.G. Leng, H. Toriumi, Y. Hamada, N. Akamatsu, S. Ohno, Ultrastructural study of upper surface layer in rat articular cartilage by “in vivo cryotechnique” combined with various treatments, Med. Electron Microsc. 33 (2000) 16–24.

[15] C.W. McCutchen, The frictional properties of animal joints, Wear 5 (1962) 1–17.

[16] A. Benninghoff, Form und Bau der Gelenkknorpel in ihren Beziehungen zur Funktion, Z. Anat. Entwicklungsgesch. 76 (1925) 43–63.

[17] U. Upadhyay, S. Kolla, L.K. Chelluri, Extracellular matrix composition analysis of human articular cartilage for the development of organ-on-a-chip, Biochem. Biophys. Res. Commun. 667 (2023) 81–88.

[18] Y. Dang, A.A. Cole, G.A. Homandberg, Comparison of the catabolic effects of fibronectin fragments in human knee and ankle cartilages, Osteoarthritis Cartilage 11 (2003) 538–547.

[19] G.A. Homandberg, F. Hui, High-Concentrations of Fibronectin Fragments Cause Short-Term Catabolic Effects in Cartilage Tissue While Lower Concentrations Cause Continuous Anabolic Effects, Arch. Biochem. Biophys. 311 (1994) 213–218.

[20] H.S. Hwang, S.J. Park, E.J. Cheon, M.H. Lee, H.A. Kim, Fibronectin fragment-induced expression of matrix metalloproteinases is mediated by MyD88-dependent TLR-2 signaling pathway in human chondrocytes, Arthritis Res. Ther. 17 (2015) 1–12.

[21] N. Burton-Wurster, G. Lust, J.N. Macleod, Cartilage fibronectin isoforms: in search of functions for a special population of matrix glycoproteins, Matrix Biol. 15 (1997) 441–454.

[22] S.A. Flowers, A. Zieba, J. Örnros, C. Jin, O. Rolfson, L.I. Björkman, T. Eisler, S. Kalamajski, M. Kamali-Moghaddam, N.G. Karlsson, Lubricin binds cartilage proteins, cartilage oligomeric matrix protein, fibronectin and collagen II at the cartilage surface, Sci. Rep. 7 (2017) 1–11.

23. K.A. Elsaid, C.O. Chichester, G.D. Jay, Lubricin Purified from Bovine Synovial Fluid and from Articular cartilage exhibit similar binding affinities to Cartilage Matrix Proteins, Transactions of the Orthopaedic Research Society 32 (2007) 551.

[24] R.C. Andresen Eguiluz, S.G. Cook, C.N. Brown, F. Wu, N.J. Pacifici, L.J. Bonassar, D. Gourdon, Fibronectin mediates enhanced wear protection of lubricin during shear, Biomacromolecules 16 (2015). 10.1021/acs.biomac.5b00810.

[25] Y. Zhu, L. Sun, Y. Wang, L. Cai, Z. Zhang, Y. Shang, Y. Zhao, A Biomimetic Human Lung-on-a-Chip with Colorful Display of Microphysiological Breath, Adv. Mater. 34 (2022) e2108972.

[26] J. Seror, Y. Merkher, N. Kampf, L. Collinson, A.J. Day, A. Maroudas, J. Klein, Normal and Shear Interactions between Hyaluronan–Aggrecan Complexes Mimicking Possible Boundary Lubricants in Articular Cartilage in Synovial Joints, Biomacromolecules 13 (2012) 3823–3832.

[27] D.P. Chang, F. Guilak, G.D. Jay, S. Zauscher, Interaction of lubricin with type II collagen surfaces: Adsorption, friction, and normal forces, J. Biomech. (2013) 1–8.

[28] D.P. Chang, N.I. Abu-Lail, F. Guilak, G.D. Jay, S. Zauscher, Conformational mechanics, adsorption, and normal force interactions of lubricin and hyaluronic acid on model surfaces, Langmuir 24 (2008) 1183–1193.

[29] S. Das, X. Banquy, B. Zappone, G.W. Greene, G.D. Jay, J.N. Israelachvili, Synergistic Interactions between Grafted Hyaluronic Acid and Lubricin Provide Enhanced Wear Protection and Lubrication, Biomacromolecules 14 (2013) 1669−1677.

[30] J. Yu, X. Banquy, G.W. Greene, D.D. Lowrey, J.N. Israelachvili, The boundary lubrication of chemically grafted and cross-linked hyaluronic acid in phosphate buffered saline and lipid solutions measured by the surface forces apparatus, Langmuir 28 (2012) 2244–2250.

[31] S.E. Majd, R. Kuijer, A. Köwitsch, T. Groth, T.A. Schmidt, P.K. Sharma, Both Hyaluronan and Collagen Type II Keep Proteoglycan 4 (Lubricin) at the Cartilage Surface in a Condition That Provides Low Friction during Boundary Lubrication, Langmuir 30 (2014) 14566– 14572.

[32] S.G. Cook, Y. Guan, N.J. Pacifici, C.N. Brown, E. Czako, M.S. Samak, L.J. Bonassar, D. Gourdon, Dynamics of Synovial Fluid Aggregation under Shear, Langmuir 35 (2019) 15887−15896.

[33] L. Hao, T. Li, L. Wang, X. Shi, Y. Fan, C. Du, Y. Wang, Mechanistic insights into the adsorption and bioactivity of fibronectin on surfaces with varying chemistries by a combination of experimental strategies and molecular simulations, Bioact. Mater. 6 (2021) 3125–3135.

[34] Z. Markovic, A. Lustig, J. Engel, Shape and Stability of Fibronectin in Solutions of Different pH and Ionic Strength, Hoppe-Seyler’s Zeitschrift Für Physiologische Chemie 364 (1983) 1795–1804.

[35] E.C. Williams, P.A. Janmey, J.D. Ferry, D.F. Mosher, Conformational states of fibronectin. Effects of pH, ionic strength, and collagen binding, J. Biol. Chem. 257 (1982) 14973–14978.

[36] B. Pardi, S.T. Ahmed, S.J. Flores, W. Flores, J.-M. Friedt, L.L.E. Mears, B.Y. Soto, R.C.A. Eguiluz, pyQCM-BraTaDio: A tool for visualization, data mining, and modelling of Quartz crystal microbalance with dissipation data, J. Open Source Softw. 9 (2024) 6831.

[37] B. Du, D. Johannsmann, Operation of the Quartz Crystal Microbalance in Liquids: Derivation of the Elastic Compliance of a Film from the Ratio of Bandwidth Shift and Frequency Shift, Langmuir 20 (2004) 2809–2812.

[38] A. Savitzky, M.J.E. Golay, Smoothing and Differentiation of Data by Simplified Least Squares Procedures, Anal. Chem. 36 (1964) 1627–1639.

39. P02751. FINC_HUMAN, UniProt (n.d.). https://www.uniprot.org/uniprotkb/P02751/entry#sequences (accessed April 7, 2025).

[40] K. Stapelfeldt, S. Stamboroski, I. Walter, N. Suter, T. Kowalik, M. Michaelis, D. Brüggemann, Controlling the multiscale structure of nanofibrous fibrinogen scaffolds for wound healing, Nano Lett. 19 (2019) 6554–6563.

[41] J. Grdadolnik, Y. Marechal, Bovine serum albumin observed by infrared spectrometry II Hydration mechanisms and interaction configurations of embedded H2O molecules, Biopolymers (Biospectroscopy) 62 (2001) 54–67. 10.1002/1097-0282(2001)62:1.

[42] A. Micsonai, É. Bulyáki, J. Kardos, BeStSel: From secondary structure analysis to protein fold prediction by circular dichroism spectroscopy, Methods Mol. Biol. 2199 (2021) 175– 189.

[43] S. Patel, A.F. Chaffotte, F. Goubard, E. Pauthe, Urea-induced sequential unfolding of fibronectin: a fluorescence spectroscopy and circular dichroism study, Biochemistry 43 (2004) 1724–1735.

[44] J.L. Hutter, J. Bechhoefer, Calibration of atomic-force microscope tips, Rev. Sci. Instrum. 64 (1993) 1868–1873.

[45] H.-J. Butt, M. Jaschke, Calculation of thermal noise in atomic force microscopy, Nanotechnology 6 (1995) 1–7.

[46] D. Nečas, P. Klapetek, Gwyddion: An open-source software for SPM data analysis, Cent. Eur. J. Phys. 10 (2012) 181–188.

[47] P. Virtanen, R. Gommers, T.E. Oliphant, M. Haberland, T. Reddy, D. Cournapeau, E. Burovski, P. Peterson, W. Weckesser, J. Bright, S.J. van der Walt, M. Brett, J. Wilson, K.J. Millman, N. Mayorov, A.R.J. Nelson, E. Jones, R. Kern, E. Larson, C.J. Carey, İ. Polat, Y. Feng, E.W. Moore, J. VanderPlas, D. Laxalde, J. Perktold, R. Cimrman, I. Henriksen, E.A. Quintero, C.R. Harris, A.M. Archibald, A.H. Ribeiro, F. Pedregosa, P. van Mulbregt, SciPy 1.0 Contributors, SciPy 1.0: fundamental algorithms for scientific computing in Python, Nat. Methods 17 (2020) 261–272.

[48] F. Lundh, An introduction to tkinter, URL: www.pythonware.com/library/tkinter/introduction/index.htm (n.d.).

[49] A.G. Hemmersam, K. Rechendorff, M. Foss, D.S. Sutherland, F. Besenbacher, Fibronectin adsorption on gold, Ti-, and Ta-oxide investigated by QCM-D and RSA modelling, J. Colloid Interface Sci. 320 (2008) 110–116.

[50] M. Bergkvist, J. Carlsson, S. Oscarsson, Surface-dependent conformations of human plasma fibronectin adsorbed to silica, mica, and hydrophobic surfaces, studied with use of Atomic Force Microscopy, Journal of Biomedical Materials Research - Part A 64 (2003) 349–356.

[51] M.D. Heath, B. Henderson, S. Perkin, Ion-specific effects on the interaction between fibronectin and negatively charged mica surfaces, Langmuir 26 (2010) 5304–5308.

[52] S. Halonen, T. Kangas, M. Haataja, U. Lassi, Urea-water-solution properties: Density, viscosity, and surface tension in an under-saturated solution, Emission Contr. Sci. Technol. 3 (2017) 161–170.

[53] K. Kawahara, C. Tanford, Viscosity and density of aqueous solutions of urea and guanidine hydrochloride, J. Biol. Chem. 241 (1966) 3228–3232.

[54] Y. Mao, J.E. Schwarzbauer, Fibronectin fibrillogenesis, a cell-mediated matrix assembly process, Matrix Biol. 24 (2005) 389–399.

[55] V. Llopis-Hernández, M. Cantini, C. González-García, Z.A. Cheng, J. Yang, P.M. Tsimbouri, A.J. García, M.J. Dalby, M. Salmerón-Sánchez, Material-driven fibronectin assembly for high-efficiency presentation of growth factors, Sci. Adv. 2 (2016) e1600188.

[56] P. Rico, J.C. Rodríguez Hernández, D. Moratal, G. Altankov, M. Monleón Pradas, M. Salmerón-Sánchez, Substrate-induced assembly of fibronectin into networks: influence of surface chemistry and effect on osteoblast adhesion, Tissue Eng. Part A 15 (2009) 3271– 3281.

[57] L.P. Kozlowski, IPC - Isoelectric Point Calculator, Biol. Direct 11 (2016) 55.

[58] D. Johannsmann, A. Langhoff, C. Leppin, I. Reviakine, A.M.C. Maan, Effect of Noise on Determining Ultrathin-Film Parameters from QCM-D Data with the Viscoelastic Model, Sensors 23 (2023). 10.3390/s23031348.

[59] W. Guo, A.A. Makarov, A.V. Buevich, Y. Jiang, Strategy for improving circular dichroism spectra deconvolution accuracy for macrocyclic peptides in drug discovery, J. Pharm. Biomed. Anal. 252 (2025) 116476.

[60] S.M. Kelly, T.J. Jess, N.C. Price, How to study proteins by circular dichroism, Biochim. Biophys. Acta 1751 (2005) 119–139.

[61] E. Oesterlund, The secondary structure of human plasma fibronecfin: conformational changesinduced by acidic pH and elevated temperatures; a circular dichroic study, Biochimica et Biophysica Acta 955 (1988) 330–336.

[62] R.W. Woody, Contributions of tryptophan side chains to the far-ultraviolet circular dichroism of proteins, Eur. Biophys. J. 23 (1994) 253262.

[63] A. Rodger, D. Marshall, Beginners guide to circular dichroism, Biochem. (Lond.) 43 (2021) 58–64.

[64] A.S. Mann, A.M. Smith, J.O. Saltzherr, A. Gopinath, R.C. Andresen Eguiluz, Glycosaminoglycans and glycoproteins influence the elastic response of synovial fluid nanofilms on model oxide surfaces, Colloids Surf. B Biointerfaces 213 (2022) 112407.

[65] J. Fan, C. Myant, R. Underwood, P. Cann, Synovial fluid lubrication of artificial joints: protein film formation and composition, Faraday Discuss. 156 (2012) 69.

[66] I.N. Serratos, A.S. Luviano, C. Millan-Pacheco, J. Morales-Corona, E.J. Alvarado Muñoz, J. Campos-Terán, R. Olayo, Quartz crystal microbalance application and in silico studies to characterize the interaction of bovine serum albumin with plasma polymerized pyrrole surfaces: Implications for the development of biomaterials, Langmuir 39 (2023) 11213– 11223.

[67] H.P. Felgueiras, N.S. Murthy, S.D. Sommerfeld, M.M. Brás, V. Migonney, J. Kohn, Competitive Adsorption of Plasma Proteins Using a Quartz Crystal Microbalance, ACS Appl. Mater. Interfaces 8 (2016) 13207–13217.

[68] A.R.C. Jones, J.P. Gleghorn, C.E. Hughes, L.J. Fitz, R. Zollner, S.D. Wainwright, B. Caterson, E.A. Morris, L.J. Bonassar, C.R. Flannery, Binding and Localization of Recombinant Lubricin to Articular Cartilage Surfaces, J. Orthop. Res. 12 (2007) 10–12.

[69] J.P. Gleghorn, L.J. Bonassar, Lubrication mode analysis of articular cartilage using Stribeck surfaces, J. Biomech. 41 (2008) 1910–1918.

[70] B. Zappone, G.W. Greene, E. Oroudjev, G.D. Jay, J.N. Israelachvili, Molecular aspects of boundary lubrication by human lubricin: effect of disulfide bonds and enzymatic digestion, Langmuir 24 (2008) 1495–1508.

[71] E.L. Radin, D.A. Swann, P.A. Weisser, Separation of a Hyaluronate-free Lubricating Fraction from Synovial Fluid, Nature 22 (1970) 377–378.

[72] M. Prajapati, K. Vishwanath, L. Huang, M. Colville, H. Reesink, M. Paszek, L.J. Bonassar, Specific degradation of the mucin domain of lubricin in synovial fluid impairs cartilage lubrication, ACS Biomater. Sci. Eng. 10 (2024) 6915–6926.

[73] J. Seror, Y. Merkher, N. Kampf, L. Collinson, A.J. Day, A. Maroudas, J. Klein, Articular cartilage proteoglycans as boundary lubricants: structure and frictional interaction of surface-attached hyaluronan and hyaluronan--aggrecan complexes, Biomacromolecules 12 (2011) 3432–3443.

[74] Y. Cao, D. Jin, N. Kampf, J. Klein, Origins of synergy in multilipid lubrication, Proc. Natl. Acad. Sci. U. S. A. 121 (2024) e2408223121.

