## Supplemental Information for "Surface-immobilized fibronectin conformation influences synovial fluid adsorption and film formation"

**AUTHOR INFORMATION**

**ORCID**

Syeda Tajin Ahmed: 0000-0002-2719-9641

Lenka Vitkova: 0000-0002-6747-1785

Warren Flores: 0000-0002-4788-4611

Ummay Honey: 0000-0002-7824-3163

Diego R. Jaramillo Pinto: 0009-0002-5615-3527

Kaleb Cutter: 0009-0004-9117-0538

Katelyn L. Lunny: 0009-0005-3565-3583

Yidan Wen: 0009-0000-3200-387X

Kevin De France: 0000-0002-5545-4793

Roberto C. Andresen Eguiluz: 0000-0002-5209-4112

**Corresponding author**

*

### Functionalization of gold-coated quartz substrates

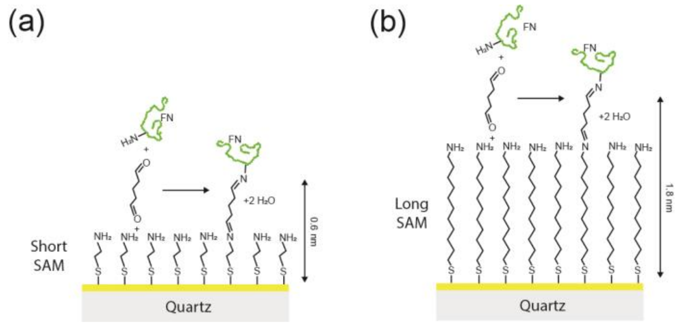


**Figure S1**. Chemical immobilization of FN to amine-functionalized surfaces with glutaraldehyde. (a) Short SAMs and (b) long SAMs.

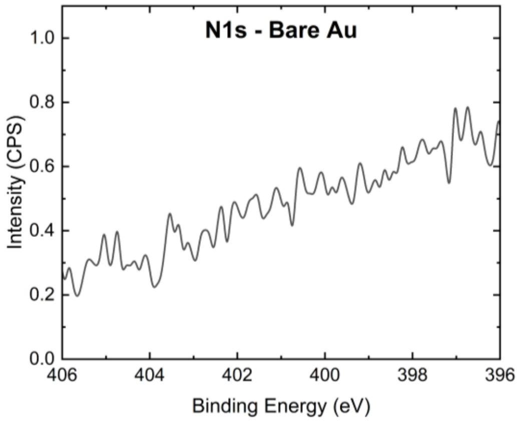


**Figure S2**. X-Ray photoelectron spectrogram for bare gold crystals, as a negative control.

### Determination of Sauerbrey mass and thin film compliance

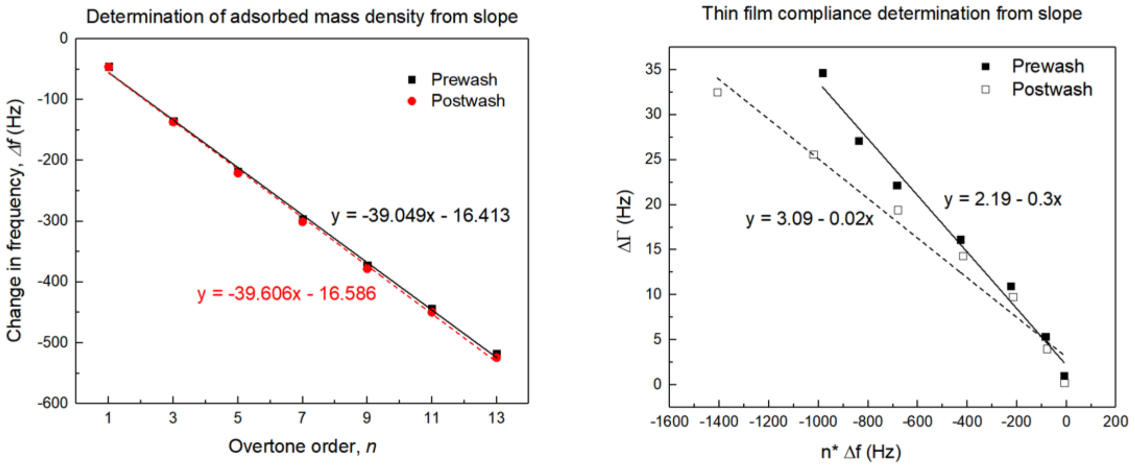


**Figure S3**. (a) For thin film adsorption in liquid, the Sauerbrey mass is estimated from the slope of change in frequency, Δf vs. overtone order, n, eq 1 from the main text. (b) Changes of half-band half width as a function of changes in frequency times number of overtone order, $n \cdot \Delta f$, are used to determine thin film compliance, $J_f'$, from the slope, eq. 2, from the main text.

### Fibronectin's concentration dependence on Sauerbrey mass and thin film compliance

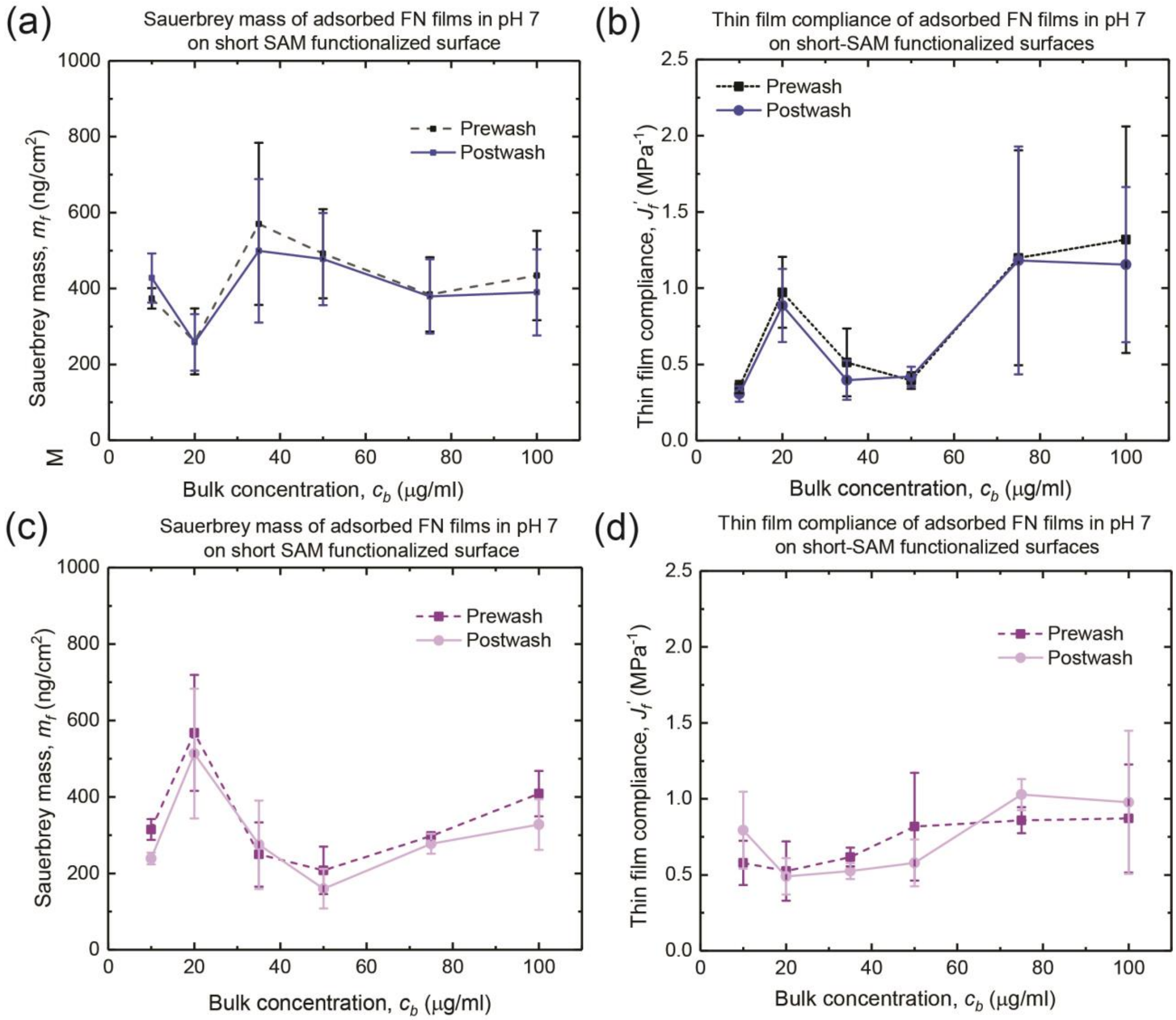


**Figure S4.** (a) The adsorption of FN films on short-SAM functionalized gold-coated crystals at pH 7 as a function of bulk concentration of FN between 10 – 100 μg/ml. (b) Thin film compliance of adsorbed FN films. (c) Sauerbrey mass of dSF films adsorbed on FN films and (d) Thin film compliance of the adsorbed dSF films on deposited FN films at pH 7. The lines are guides for eye.

**Diffuse Reflectance Circular Dichroism (DRCD) data unit conversion**

The unit conversion was performed to obtain spectra in concentration-independent units. We used molar residue ellipticity (*MRE*), as this unit is the standard for representing liquid sample CD data. For liquid (bulk) samples:

$$MRE_{liq} = \frac{\theta}{c \cdot l \cdot N_r} \qquad \text{eq. S1}$$

where $\theta$ is the measured ellipticity [mdeg], $c$ is the protein molar concentration [mol·dm$^{-3}$], l is the path length [cm], and $N_r$ is the number of amino acid residues. For FN, N is 4956 residues. Analogously, the molar concentration of protein adsorbed on the substrate was determined as follows:

$$c = \frac{c_{surf}}{t} \qquad \text{eq. S2}$$

where $c_{surf}$ [mol·m$^2$] is the molar surface concentration and $t$ [m] is the thickness of the film. To determine the molar surface concentration, the following argument was formulated:

$$c_{surf} = \frac{n}{A} = \frac{m}{M_w \cdot A} \qquad \text{eq. S3}$$

where $n$ [mol] is the amount of protein and $A$ [m$^2$] is the sensor area. The amount of the adsorbed substance can be found as the mass of the substance $m$ [g] divided by its molar mass $M_w$ [g·mol$^{-1}$].

To determine the mass of adsorbed protein, Sauerbrey mass $m_{Sauerbrey}$ [µg·mol$^{-1}$] was used based on the following assumption:

$$\Delta m \equiv m \qquad \text{eq. S4}$$

$$m_{Sauerbrey} = \frac{\Delta m}{A} \qquad \text{eq. S5}$$

making the molar concentration of adsorbed protein:

$$c = \frac{m_{Sauerbrey}}{M_w \cdot t} \qquad \text{eq. S6}$$

This assumption leads to overestimation of the protein concentration, as it disregards the solvent's contribution to $m_{Sauerbrey}$. Nevertheless, we believe that this method is accurate for resolving the concentration dependence of the measured ellipticity.

The film thickness can be assigned to Sauerbrey thickness $d_{Sauerbrey}$ [nm], thus:

$$c = \frac{m_{Sauerbrey}}{M_w \cdot d_{Sauerbrey}} \qquad \text{eq. S7}$$

The path length is determined by the path that the light travels through the sample, which is the film thickness, earlier assigned as $d_{Sauerbrey}$. Due to the reflection mode setup of the experiment, the light travels through the adsorbed film twice, making the path length:

$$l = 2 \cdot d_{Sauerbrey} \qquad \text{eq. S8}$$

With these assumptions, the *MRE* conversion formula can be written as:

$$MRE_{film} = \frac{\theta}{\frac{m_{Sauerbrey}}{M_W \cdot d_{Sauerbrey}} \cdot 2 \cdot d_{Sauerbrey} \cdot N_r} \quad \text{eq. S9}$$

or after simplification:

$$MRE_{film} = \frac{\theta \cdot M_W}{2 \cdot m_{Sauerbrey} \cdot N_r} \quad \text{eq. S10}$$

This formula yields $[MRE_{film}]$ = mdeg·m$^2$·mol$^{-1}$, which is equal to the units of standard *MRE* converted liquid samples. While the assumption of $m_{\text{Sauerbrey}}$ leads to overestimation, we maintain the value of this conversion for qualitative comparison of CD spectra of protein films in both terms of peak position and amplitude.

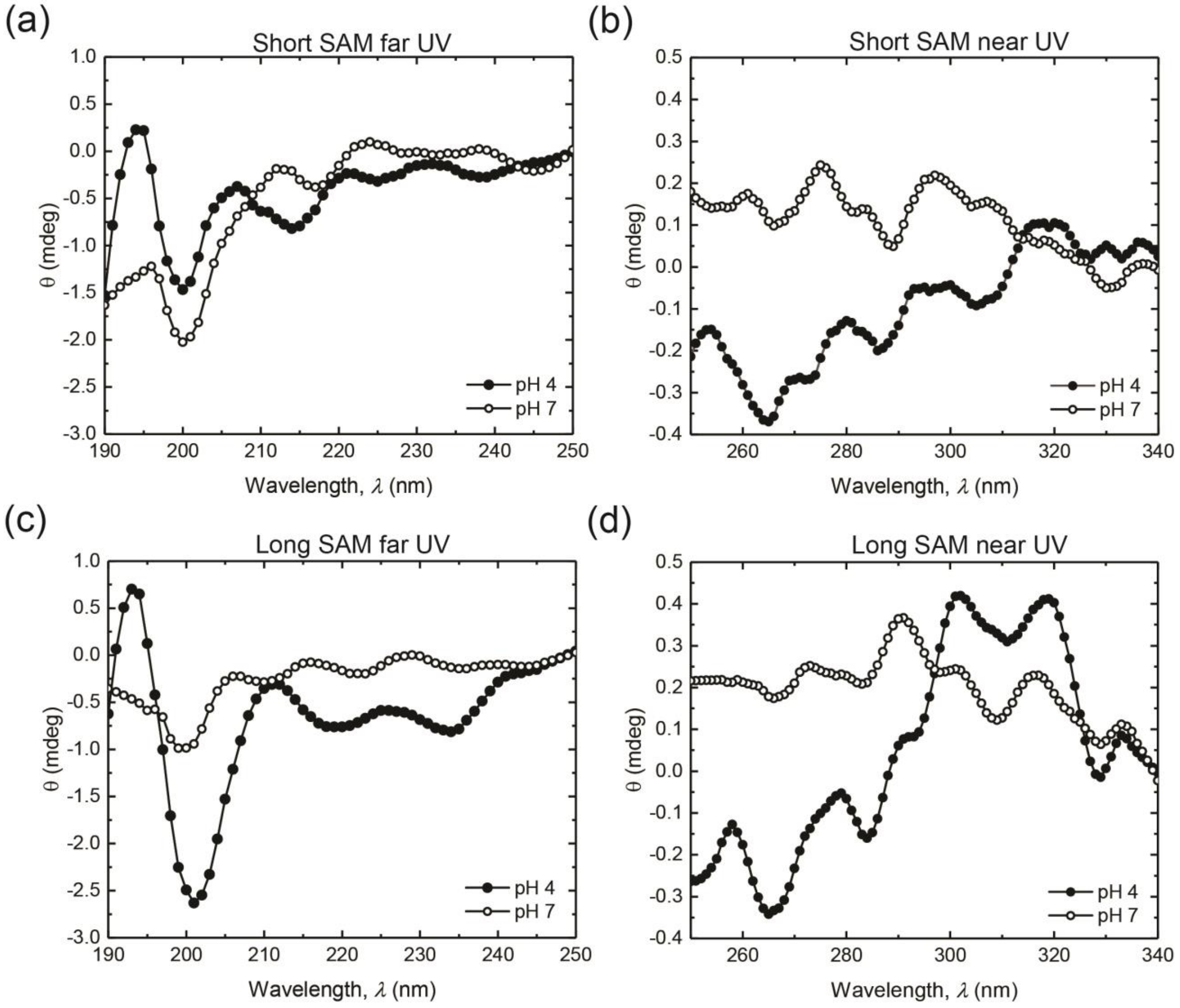


**Figure S5.** (a) Far UV DRCD raw spectra of FN adsorbed onto a short SAM at pH 4 and pH 7, (b) far UV DRCD raw spectra of FN adsorbed onto a long SAM at pH 4 and pH 7, (c) near UV DRCD raw spectra of FN adsorbed onto short SAM at pH 4 and pH 7, and (d) near UV DRCD raw spectra of FN adsorbed onto a long SAM at pH 4 and pH 7.

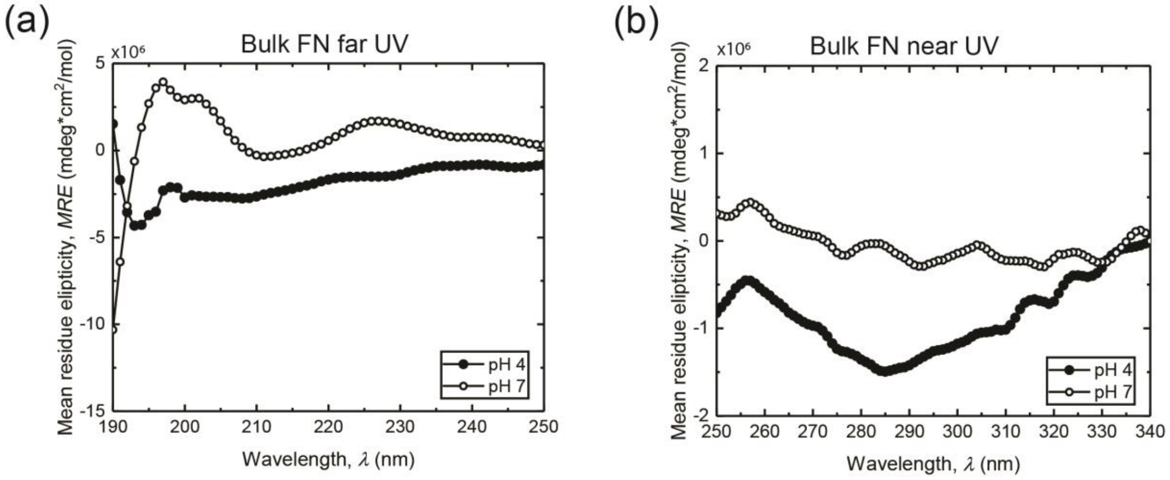


**Figure S6.** Circular dichroism of fibronectin (FN) at a bulk concentration of 50 μg/mL in PBS at pH 7 and at pH 4. (a) Far UV CD spectra of FN at pH 4 and pH 7and (b) near UV CD spectra of FN at pH 4 and pH 7.

### Aqueous buffer pH switch baseline drift

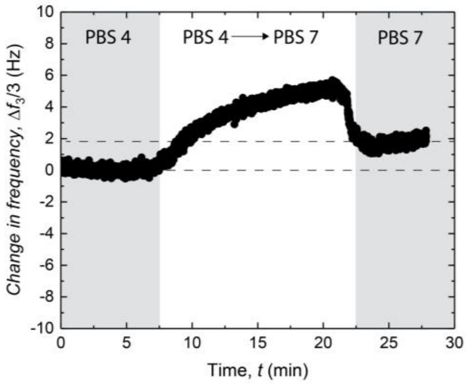


**Figure S7.** Representative curve, showing a baseline shift due to buffer switch from pH 4 to pH 7.

**Atomic force microscopy film morphologies of control samples and films in the height channel**

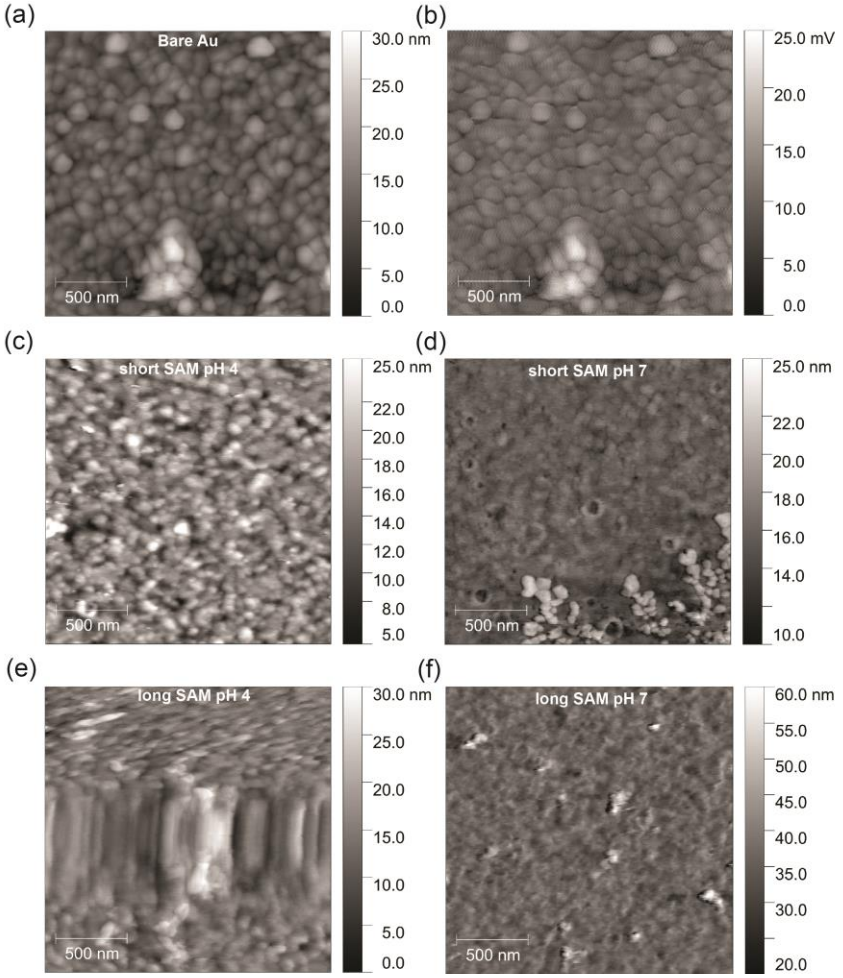


**Figure S8.** Representative AFM images of (a) height signal obtained for a bare Au on a quartz crystal used for QCM-D adsorption measurements, (b) and its corresponding friction signal. (c) FN on short-SAM functionalized gold-coated crystals deposited at pH 4, (d) FN on short-SAM functionalized gold-coated crystals deposited at pH 7, (e) FN on long-SAM functionalized gold-coated crystals deposited at pH 4, and (f) FN on long-SAM functionalized gold-coated crystals deposited at pH 7.

### QCM-D control adsorption measurements of dSF onto various surfaces

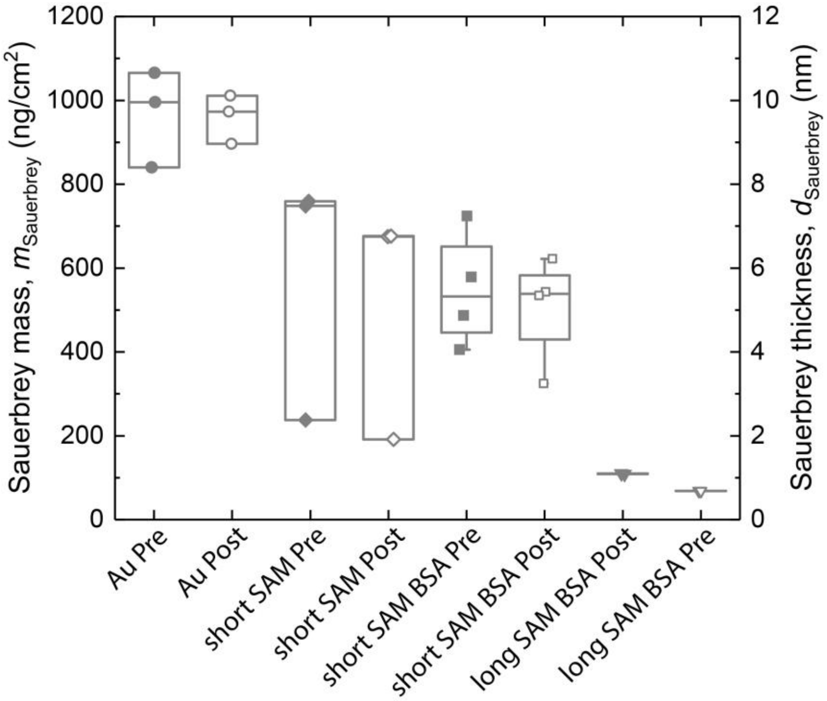


**Figure S9.** Sauerbrey mass, $m_{Sauerbrey}$ of dSF adsorbed onto various control surfaces. These are: clean Au surfaces, before (pre) and after (post) PBS 7 wash, short SAM functionalized Au surfaces before (pre) and after (post) PBS 7 wash, and short SAM functionalized Au surfaces previously blocked with a BSA step, before (pre) and after (post) PBS 7 wash.

**Table S1.** Summary of $m_{Sauerbrey}$ and $J'_f$ of dSF films formed on various control surfaces

| | **Pre-wash $m_{Sauerbrey}$ (ng/cm$^2$)** | **Post-wash $m_{Sauerbrey}$ (ng/cm$^2$)** | **Pre-wash $J'_f$ $\cdot 10^{-7}$ (Pa$^{-1}$)** | **Post-wash $J'_f$ $\cdot 10^{-7}$ (Pa$^{-1}$)** |
|---|---|---|---|---|
| On short SAM | 600 ± 200 | 500 ± 200 | 7 ± 2 | 7 ± 1 |
| With BSA on short SAM | 580 ± 60 | 530 ± 50 | 6 ± 1 | 5 ± 1 |
| With BSA on long SAM | 109 ± 1 | 69 ± 1 | 22 ± 1 | 24± 1 |
| On bare Au | 970 ± 70 | 960 ± 30 | 5 ± 1 | 5 ± 1 |

### QCM-D control adsorption measurements of BSA onto various surfaces

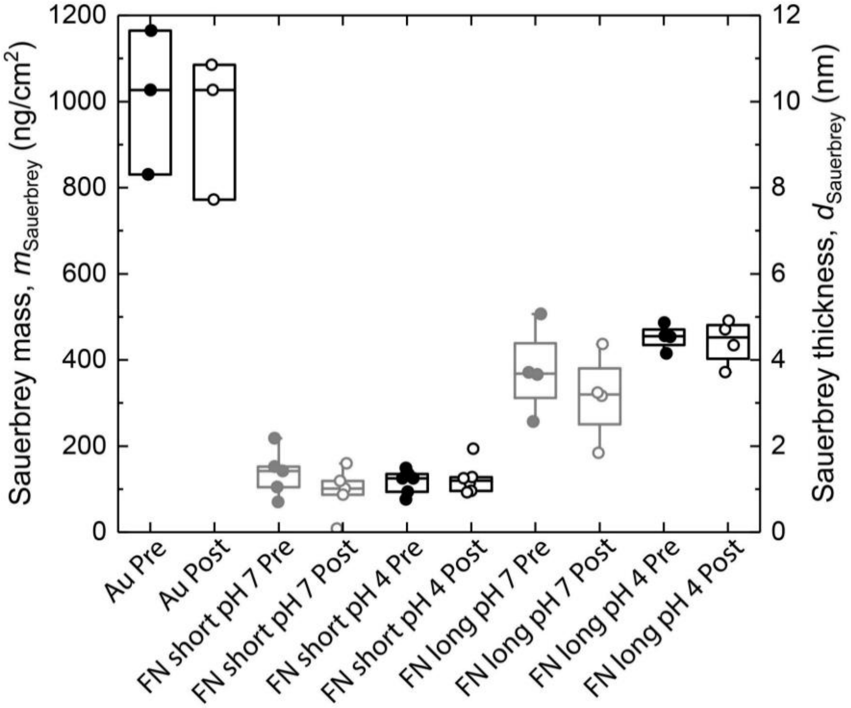


**Figure S10.** Sauerbrey mass, $m_{Sauerbrey}$, of BSA adsorbed onto various control surfaces These are: clean Au surfaces, before (pre) and after (post) PBS at pH 7 wash, FN on short SAM functionalized Au surfaces before (pre) and after (post) PBS 7 wash, FN on short SAM functionalized Au surfaces before (pre) and after (post) PBS 4 wash, FN on long SAM functionalized Au surfaces before (pre) and after (post) PBS 7 wash, and FN on long SAM functionalized Au surfaces before (pre) and after (post) PBS 4.

**Table S*2*.** Summary of $m_{Sauerbrey}$ of BSA films formed on FN film before (pre) and after (post) PBS washes on short and long SAMs.

| SAM - pH | Pre-wash $m_{Sauerbrey}$ (ng/cm$^2$) | Post-wash $m_{Sauerbrey}$ (ng/cm$^2$) | Pre-wash $J'_f \cdot 10^{-7}$ (Pa$^{-1}$) | Post-wash $J'_f \cdot 10^{-7}$ (Pa$^{-1}$) |
|---|---|---|---|---|
| Short - 7 | 137 ± 25 | 95 ± 25 | 9 ± 80 | 1 ± 7 |
| Short - 4 | 117 ± 11 | 125 ± 15 | 7 ± 2 | 7 ± 2 |
| Long - 7 | 375 ± 51 | 315 ± 52 | 7 ± 39 | 7 ± 40 |
| Long - 4 | 452 ± 15 | 442 ± 26 | 6 ± 1 | 6 ± 2 |
| On bare Au (no SAMs) | 1007 ± 97 | 961 ± 96 | 5 ± 1 | 4 ± 2 |
